# Disruption of Epithelial Integrity Drives Mesendoderm Differentiation in Human Induced Pluripotent Stem Cells by Enabling BMP and ACTIVIN Sensing

**DOI:** 10.1101/2021.10.22.465459

**Authors:** Diane Rattier, Thomas Legier, Thomas Vannier, Flavio Maina, Rosanna Dono

**Author notes:** **Correspondence to:** Rosanna Dono, Ph.D., Aix Marseille Univ, CNRS, IBDM, Turing Center for Living Systems, 163 Avenue de Luminy, case 907 - 13009 Marseille (France). These authors contributed equally: Diane Rattier, Thomas Legier.

## Abstract

The processes of primitive streak formation and fate specification in the mammalian epiblast rely on complex interactions between morphogens and tissue organization. Little is known about how these instructive cues functionally interact to regulate gastrulation. We interrogated the interplay between tissue organization and morphogens by using human induced pluripotent stem cells (hiPSCs) down-regulated for the morphogen regulator GLYPICAN-4, in which defects in tight junctions result in areas of disrupted epithelial integrity. Remarkably, this phenotype does not affect hiPSC stemness, but impacts on cell fate acquisition processes. Strikingly, cells within disrupted areas become competent to perceive BMP4 and ACTIVIN A/NODAL gastrulation signals and thus, differentiate into mesendoderm. Yet, disruption of epithelial integrity prolongs temporal activation of BMP4 and ACTIVIN A/NODAL downstream effectors and correlates with enhanced hiPSC endoderm/mesoderm differentiation potential. Altogether, our results disclose epithelial cell integrity as a key determinant of TGF-**β** activity and highlight a new mechanism guiding morphogen sensing and spatial cell fate change within an epithelium.

## INTRODUCTION

Gastrulation is a major morphogenetic process in animal development during which the embryonic germ layers are generated and the basic body plan is laid down. In mammals, gastrulation is first marked by the appearance of the primitive streak (PS), a transient structure arising by thickening of the epiblast. Epiblast cells recruited and ingressing into the primitive streak will acquire a mesendoderm (MES) fate, marked by co-expression of the transcription factors T/Brachyury (BRACH) and Eomesodermin (EOMES). Subsequently cells undergo epithelial-mesenchymal transition (EMT) to initiate differentiation into the mesoderm (ME) and definitive endoderm (DE) germ layers^1, 2^. Instead, epiblast cells that do not ingress through the streak will become the future ectodermal layer.

A mechanistic understanding of how mammalian gastrulation is initiated and how pluripotent cells of the epiblast acquire different fates has attracted considerable scientific interest given the impact that findings can have for developmental biology and for clinical application. Great insights have come from genetic and embryological studies carried out in the mouse embryo. These studies have led to the proposal that gastrulation is initiated by means of biochemical signals like WNT, the TGF-β superfamily members, BMP and ACTIVIN A/NODAL, and their inhibitors, which establish a signaling crosstalk between embryonic and extra-embryonic cells^2^. In this scenario, gastrulation is initiated when epiblast cells are instructed by BMP4, secreted by the extra embryonic ectoderm, to produce the secreted signaling protein WNT3A^2, 3^. WNT3A in turn stimulate epiblast cells to express and release NODAL that will feedback to the extraembryonic ectoderm to maintain BMP signaling in the extraembryonic ectoderm as well as the overall signaling crosstalk between the two cellular compatments^2–5^. As a result of this signaling crosstalk between BMP, WNT and NODAL, the primitive streak is formed and the epiblast becomes patterned.

Recently, this configuration has been challenged by studies showing that these signaling interactions may be further modulated by topological cues and by the epithelial organization of the epiblast. In particular, the use of sophisticated live imaging tools in mouse embryos has highlighted that BMP receptors localize at the basolateral membrane of mouse epiblast in vivo facing a narrow interstitial space located between the epiblast cells and the underlying visceral endoderm. Interestingly, the lack of tight junctions (TJs) between the extraembryonic ectoderm and the edge of the epiblast at the posterior embryo site enables the diffusion of BMP4 through this interstitial space, allowing ligands to reach basolateral localized receptors and initiate gastrulation^6^.

As these mechanical studies are arduous given the small size and inaccessibility of early post implantation embryos, human pluripotent stem cells (hPSCs) have become a reference system to dissect physical processes and the feedback interactions with biochemical signals. Pioneering studies with hPSCs grown on a disk-shaped plate, to mimic the mechanical constrains of the human epiblast (called ‘micropattern’ in the following), has highlighted physical cues, such as colony size, shape and cell density, can impact on the rate and trajectory of hPSC differentiation by acting on the balance between differentiation-inducing and -inhibiting factors^7, 8^. Subsequent experiments in which micropattern technology was applied to the study of tissue patterning have revealed a remarkable capability of hPSCs to recapitulate aspects of germ layer patterning such as simultaneous formation of the embryonic germ layers radially organized when exposed to the gastrulation initiating signal, BMP4^9, 10^. Of note, additional mechanistic studies have revealed that this in vitro patterning process relies on the epithelial architecture of the hPSC layer, which determines accessibility of BMP4 receptor in cells and formation of morphogen gradients ^11, 12^. By using this confinement strategy, it has been also proposed that regions of high cell-cell tension in the hPSC layer foster the appearance of gastrulation like nodes (PS-like) and mesoderm differentiation in response to BMP4 stimulation^13^. This is due to the release of β-catenin from cell-cell adhesion sites and its nuclear translocation. Collectively, these overall results raise the possibility that intrinsic changes in the epiblast cell layer could trigger the onset of gastrulation and MES differentiation by sensitizing cells to morphogens.

The heparan sulphate proteoglycan GLYPICAN-4 (GPC4)^14^, is a key regulator of signals controlling the balance between PSC self-renewal and differentiation (e.g. BMP, WNT, FGF)^15–20^. By performing functional analysis of human induced PSCs (hiPSCs) with reduced GPC4 protein levels (GPC4sh hiPSCs), we discovered that these cells provide a unique biological system to obtain new insights into the interplay between the physical properties of epiblast cells and biochemical signals during gastrulation.

Here we show that in contrast to control cells, GPC4sh hiPSCs display altered epithelial integrity with areas of abnormal TJs distribution. Of note, GPC4sh hiPSCs display enhanced differentiation potential into MES and ME, DE lineages, under classical differentiation protocols compared to control cells. Strikingly, areas of disrupted epithelial integrity correlate precisely with expression of the MES markers BRACH and EOMES, indicating that the morphological phenotype of the GPC4sh hiPSC layer sensitize cells to respond to morphogens. By performing stimulation assays with BMP4, ACTIVIN A and WNT3A ligands, we show that epithelial integrity regulates the ability of hiPSCs to respond to BMP4 and ACTIVIN A but not WNT3A. In addition, we demonstrate that disruption of epithelial integrity prolongs temporal activation of TGF-β signaling pathway. Thus, our findings highlight alteration of epithelial integrity as a novel mechanism for controlling TGFβ signaling and differentiation processes.

## RESULTS

### Down-regulation of GPC4 in hiPSC affects epithelial integrity by altering TJ organization

To investigate the consequences of disrupting epithelial organization following GPC4 down-regulation in hiPSCs, we used two previously published hiPSC lines in which GPC4 was down-regulated in the parental hiPSC 029 by means of two different GPC4-targeting shRNAs (cloned hiPSCs: GPC4sh5 and GPC4sh2; TableS1)^14^. Specifically, we employed the GPC4sh5-c10 and the GPC4sh2-c3 clones as representative cells carrying the GPC4sh5 and the GPC4sh2 sequences, respectively (Fig. 1). The 029 hiPSC line transduced with the non-mammalian sequence GFP was used as control (CRTLsh)^14^. RT-qPCR analysis of GPC4 expression showed no significant differences in *GPC4* transcript levels between WT and CTRLsh hiPSCs (Fig. 1a) as previously reported^14^. In contrast, *GPC4* transcript levels were reduced of 60±14% in GPC4sh5-c10 and of 70±4% in GPC4sh2-c3 hiPSCs (Fig. 1a). Consistently, immunocytochemical analysis revealed that GPC4sh5 and GPC4sh2 cultures display reduced GPC4 levels compared to WT and CTRLsh (Fig. 1b).

**Figure 1.**
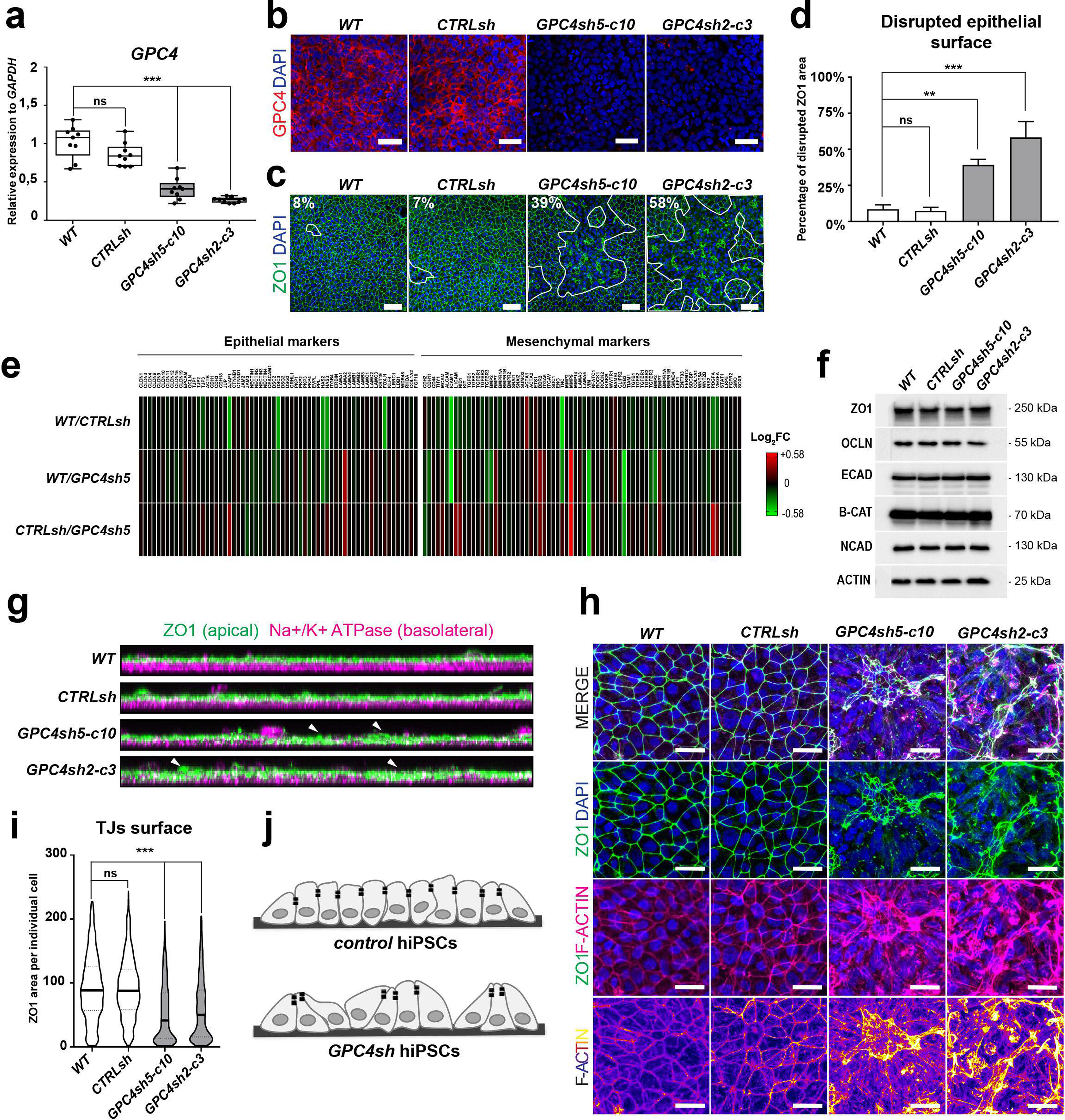
Down-regulation of GPC4 in hiPSC affects epithelial integrity by altering TJ distribution. **(a)** RT-qPCR analyses of transcript levels of *GPC4* in WT, CTRLsh, GPC4sh5-c10 and GPC4sh2-c3 029 hiPSCs showing a down-regulation of *GPC4* expression in GPC4sh hiPSCs. Transcript levels were normalized to the mean of WT ΔCt. Box plots represent the median with interquartile range, the errors bars indicate min and max values, n = 9. **(b)** Immunofluorescence analysis of GPC4 (red) and DAPI positive nuclei (blue) in WT, CTRLsh, GPC4sh5-c10 and GPC4sh2-c3 029 hiPSCs. Scale bar: 50 μm. **(c)** Immunofluorescence analysis of ZO1 (green) in WT, CTRLsh, GPC4sh5-c10 and GPC4sh2-c3 029 hiPSCs. White dashed areas highlight zones of disturbed TJs in each hiPSC lines. Scale bar: 50 μm. **(d)** The extent of TJ disruption was quantified from staining shown in (c). Data are represented as mean ± SEM, n = 9. **(e)** RNA-seq analyses of transcript levels of epithelial and mesenchymal genes in WT, CTRLsh, GPC4sh5-c10 029 hiPSCs. Epithelial and mesenchymal related genes are indicated on the figure. Data are represented as heatmap of Log2FC. **(f)** Western-blot analysis of ZO1, OCLN, ECAD, β-CAT and N-CAD in WT, CTRLsh, GPC4sh5-c10 and GPC4sh2-c3 029 hiPSC total protein extracts. Note similar protein expression level in all hiPSC lines. ACTIN protein levels were used as loading control. **(g)** Immunofluorescence analysis of ZO1 (green) and Na^+^/K^+^ ATPase (magenta), in WT, CTRLsh, GPC4sh5-c10 and GPC4sh2-c3 029 hiPSCs. Pictures are presented as a lateral view of stained cells. Arrow heads point to areas of abnormal apico-basal polarity **(h)** Immunofluorescence analysis of ZO1 (green), F-ACTIN (magenta) in WT, CTRLsh, GPC4sh5-c10 and GPC4sh2-c3 029 hiPSCs. Scale bar: 25 μm. **(i)** The surface delimited by the ZO1 protein in each individual cell was quantified from staining shown in (h). Data are represented as violin plot (distribution around the median), n= 4000 cells per hiPSC line. (**j**) Schematic representation of epithelial cell morphology of CTRLsh, and of disrupted TJs in GPC4sh5-c10. Statistical analysis for the overall figure: (d, i) one-way ANOVA followed by Dunnett’s multiple comparison test. P values: (***) < 0.001, (**) < 0.01, (*) < 0.05.

Next, we evaluated the morphology and organization of controls and GPC4sh hiPSC cultures by analysing epithelial features. In particular, cells grown at confluence were immunostained for the TJ proteins, Zona occludens 1 (ZO1) and Occludin (OCLN), by immunocytochemistry. Intriguingly, we found that GPC4sh cultures display areas with disrupted TJ organization, whereas control cultures form the classical epithelial sheet with organized and defined TJs, (Fig. 1c and Extended Data Fig. 1a). Quantification revealed that this phenotype covers 39,1% and 58,2% of the cell layer in GPC4sh2-c3 and GPC4sh5-c10 hiPSCs, respectively, whereas TJ disrupted areas account only for 8,4% and 7,4% of the epithelial sheet in control hiPSCs (Fig. 1c-d). HiPSC cultures with down-regulated GPC4 exhibit abnormal TJ organization even when cultured at low density (0,2×10^5^ cells/cm^2^ instead of 1×10^5^ cells/cm^2^), Extended Data Fig. 1b. Finally, disruption of TJ organization in GPC4sh hiPSCs occurs in cells plated on different extracellular matrix (ECM) such as Matrigel (mixture of Collagen, Laminin and growth factors), Laminin or Vitronectin (Extended Data Fig. 1c).

To assess the effect of GPC4 down-regulation in a different hiPSC line, we examined TJ organization in the previously published AICS-0023 hiPSCs targeted with CTRLsh and GPC4sh (Extended Data Fig. 1d)^14^. Analysis of ZO-1 distribution revealed extensive areas of disrupted TJ organization in the epithelial sheet of the AICS-0023 GPC4sh compared with the CTRLsh line (Extended Data Fig. 1e). This result highlights that disrupted TJ organization is triggered by GPC4 down-regulation and is not limited to one hiPSC line.

It is well known that loss of TJs occurs when epithelial cells switch to a mesenchymal state^21, 22^. Therefore, we next examined the expression of genes regulating epithelial and mesenchymal states in controls and GPC4sh hiPSCs using RNA-sequencing (RNA-seq) data. By comparing transcript levels of 62 epithelial and 71 mesenchymal markers we did not find major differences between WT, CTRLsh and sh5GPC4 hiPSCs (Fig. 1e and Supplementary Table 1). These results were corroborated by western-blot analyses of some epithelial (ZO1, OCLN, E-Cadherin (E-CAD) and β-Catenin (β-cat)) and the mesenchymal (N-Cadherin (N-CAD)) markers. Consistently with our RNA-seq data, we found similar levels of ZO1, OCLN, E-CAD, β-cat and N-CAD proteins in controls and GPC4sh hiPSCs (Fig. 1f). Thus, this overall analysis indicates that down-regulation of GPC4 in hiPSCs does not promote an epithelial to mesenchymal transition (EMT).

We then asked whether down-regulation of GPC4 disrupts other aspects of the apico-basal polarity of epithelial cells. We examined the distribution in the Z-axis of baso-lateral (Na+/K+ ATPase and E-CAD) and apical (ZO1) marker proteins^23–25^. Immunocytochemical analyses of control hiPSC cultures showed that Na+/K+ ATPase and E-CAD maintained their basolateral localization below TJs (ZO1 immunostaining). Instead, Na+/K+ ATPase or E-CAD staining were found intermingled with ZO1 in GPC4sh hiPSC cultures. These results highlighting disruption of the apico-basal cell polarity in GPC4sh hiPSCs, which can lead to an apical exposure of the baso-lateral localized proteins (Fig. 1g and Extended Data Fig. 1f).

To further characterize the changes in epithelial morphology characterizing GPC4sh hiPSCs, we analyzed the state of TJs in conjunction with that of F-ACTIN cortex. In control hiPSC cultures, we observed that the F-ACTIN cortex is clearly defined and organized in a regular pattern matching the TJ localization (Figure 1h), reflecting an equilibrated tissue tension ^26^. Instead, in GPC4sh cultures we noticed that, within the areas of altered TJs, the F-ACTIN cortex is poorly defined although displaying multiple foci of accumulation (Figure 1h), indicative of a high tissue tension^26^. Quantification of the apical cell surface size, revealed that this surface is strongly reduced in GPC4sh cultures compared to control hiPSCs (Fig. 1i-j).

Taken together, these results show that down-regulation of GPC4 in hiPSCs alters TJ distribution and apico-basal cell polarity. Although this phenotype does not affect epithelial cell identity, it disrupts the morphology and integrity of the hiPSC epithelial layer and enables apical exposure of basolateral localized proteins.

### Disruption of the epithelial integrity does not affect hiPSC stemness properties

We next asked whether disruption of the epithelial integrity in hiPSC impacts on their stemness and differentiation properties. We found that GPC4sh hiPSCs stained intensely for the stemness marker Alkaline Phosphatase as controls (Extended Data Fig. 2-1a), and that the expression levels of the Epiblast marker OTX2 and of the core pluripotency genes OCT4, NANOG and SOX2 were not affected (Extended Data Fig. 2-1b-d). Similarly, no differences were found on the cell cycle progression (Extended Data Fig. 2-1e), on the mitotic and apoptotic rates (Extended Data Fig. 2-1f-h)^14^. Results are consistent with data we reported previously^14^. To extend this analysis further, we used our RNA-seq data to follow the expression levels of 48 markers of PSCs in WT, CTRLsh and GPC4sh5 cultures maintained in self-renewal conditions. As above, the transcript levels of all PSC markers analyzed were not significantly changed between WT, CTRLsh and GPC4sh5 hiPSCs (Fig. 2a; Supplementary Table 2). Collectively, these findings show that disruption of epithelial integrity in hiPSCs does not affect stemness identity when cells are cultured in self-renewal conditions.

**Figure 2.**
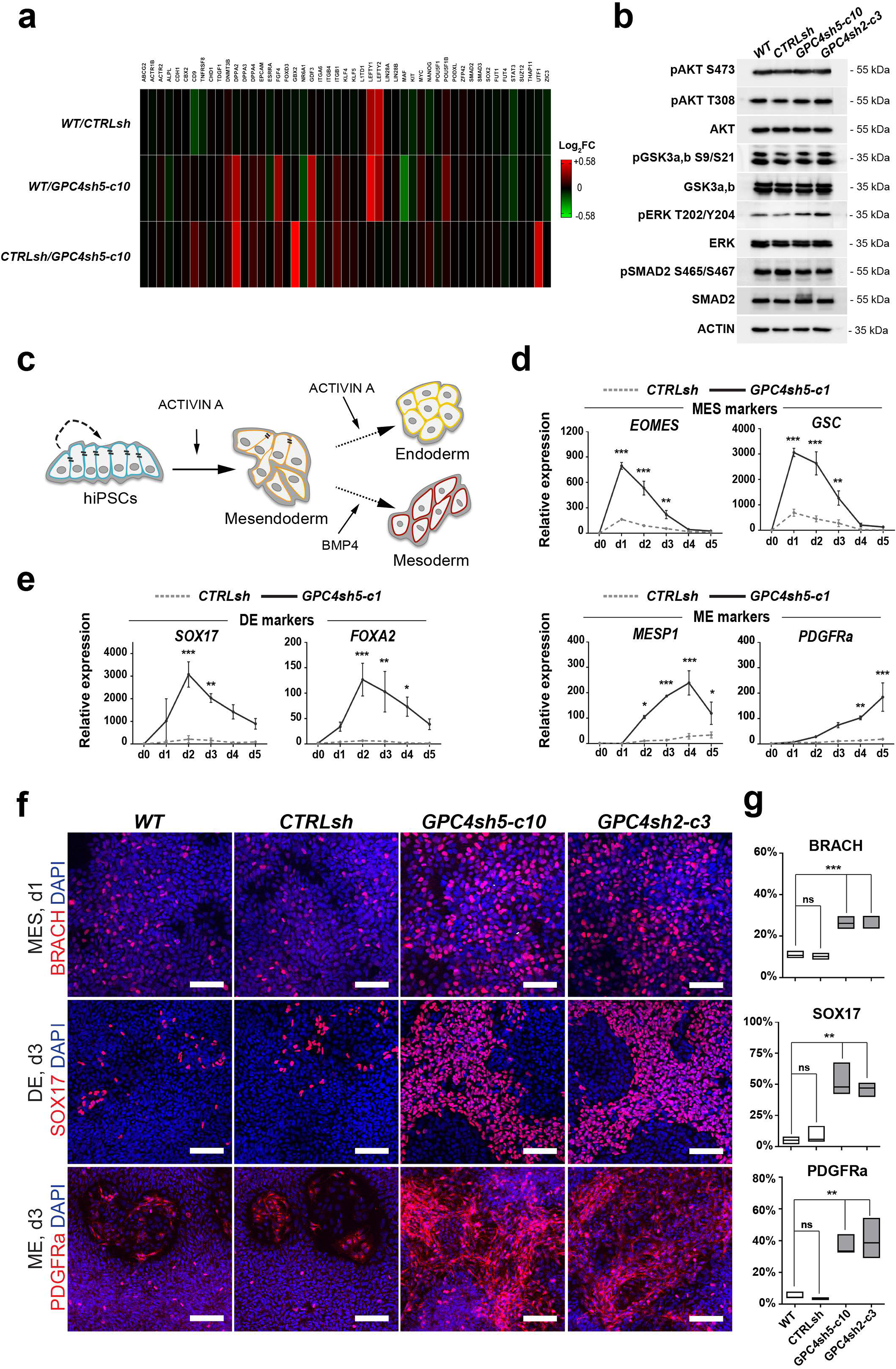
Disruption of epithelial integrity does not affect stemness properties of hiPSCs but enhances differentiation potential. (**a)** Transcripts levels of genes regulating pluripotency were analyzed by RNA-seq in WT, CTRLsh and GPC4sh5-c10 029 hiPSCs at the undifferentiated stage. Data are represented as heatmap of Log2FC. **(b)** Total protein extract of WT, CTRLsh, GPC4sh5-c10 and GPC4sh2-c3 029 hiPSCs at the undifferentiated stage were analyzed by Western-blot with pAKT S473, pAKT T308, AKT, pERK T202/Y204, ERK, pGSK3a, b S9/S21, GSK3a, b, pSMAD2 S465/S467 and SMAD2 antibodies. Note similar expression level and phosphorylation of AKT, ERK, GSK3a, b and SMAD2 proteins in all hiPSC lines. ACTIN protein levels were used as loading control. **(c)** Scheme depicting the strategy of differentiation into MES and subsequently into DE and ME used. **(d, e)** CTRLsh and GPC4sh5-c10 029 hiPSCs were differentiated for 5 days into DE or ME and transcript levels of MES, DE or ME specific markers were analyzed by RT-qPCR. (d): MES markers (*EOMES* and *GSC*), (e): DE markers (*SOX17* and *FOXA2*), (e): ME markers (*MESP1* and *PDGFRa*). Data were normalized to the d0 of CTRLsh and represented as mean ± SEM, n = 3. **(f)** Immunofluorescence analysis of BRACH (red, MES), SOX17 (red, DE) or PDGFRa (red, ME) on WT, CTRLsh, GPC4sh5-c10 and GPC4sh2-c3 029 hiPSCs differentiated for 1 day into MES, 3 days into DE or 3 days into ME. Scale bar: 100 μm. **(g)** Percentages of BRACH, SOX17 and PGFDRa positive cells in WT, CTRLsh, GPC4sh5-c10 and GPC4sh2-c3 029 hiPSCs were quantified from staining shown in (f). Box plots represent the median with min and max values, n = 3. Statistical analysis for the overall figure: (d, e) two-way ANOVA followed by Sidak’s multiple comparison test. (g) one-way ANOVA, followed by Dunnett’s multiple comparison test. P values: (***) < 0.001, (**) < 0.01, (*) < 0.05.

Previous studies have shown that the balance between self-renewal and differentiation of PSCs is coordinated by the crosstalk between the PI3K/AKT, RAF/MEK/ERK, and WNT/GSK3b signaling pathways that together impact on the ability of ACTIVIN A/SMAD2,3 to promote either self-renewal or differentiation into MES cells^27–31^. Therefore, we determined whether disruption of hiPSC epithelial integrity impacts on this regulatory signaling network. Western blot analysis showed similar activation levels of AKT, ERK, GSK3a/b and SMAD2 pathways in control and GPC4sh hiPSCs maintained in self-renewal conditions (Fig. 2b). Consistently, RNA-seq analysis revealed that genes regulating fate acquisition such as *GSC*, *BRACH*, *EOMES*, *FOXC1*, *FOXA1* and *SOX1* were not expressed neither in WT and CTRLsh hiPSCs nor in GPC4sh5 cells. Together these results show that disruption of hiPSC epithelial integrity does not impact on self-renewal and pluripotency of hiPSCs neither primes them to express cell lineage markers when cells are cultured under stemness conditions.

### Enhanced Mesendoderm fate acquisition in GPC4sh cultures

Recent studies have shown that differentiation of hiPSCs towards the MES lineage and into its DE and ME derivatives rely on hiPSC mechanics and epithelial organization properties^10, 11, 13, 32^. Given the peculiar morphological phenotype of GPC4sh hiPSCs, we reasoned that GPC4sh cultures could be a powerful system to study the impact of epithelial integrity on hiPSC differentiation along MES, DE and ME embryonic lineages. HiPSCs were exposed to ACTIVIN A, a surrogate for NODAL in hPSC cultures, and BMP4 (Fig. 2c), which trigger differentiation along these lineages both in vitro and in embryos^33, 34^. We then evaluated the differentiation profile of control and GPC4sh hiPSCs by using RT-qPCR analysis following lineage specific markers.

In agreement with published studies, the MES markers *EOMES*, *GSC*, *NODAL* and *MIXL1* were already expressed at day 1 (d1) of differentiation in control cells exposed to ACTIVIN A, and transcript levels were gradually down-regulated from d1 to d5 (Fig. 2d and Extended Data Fig. 2-2a-b). When cells were kept in the presence of ACTIVIN A, expression of the DE markers *SOX17*, *FOXA2*, *CER1* and *CXCR4* started at d1, peaked between d2 and d3 to gradually reduced from d3 to d5 (Fig. 2e and Extended Data Fig. 2-2a-b)^33, 35, 36^. Instead, when ACTIVIN A was replaced by BMP4 at d2, the early ME markers *MESP1* and *TBX6* were gradually expressed from d2 to d4 then down-regulated from d4/d5 (Fig. 2e and Extended Data Fig. 2-2a-b). At these late time points of differentiation, we observed concomitant expression of late ME markers, such as *VEGFR2* and *PDGFRa*^34, 37^ (Fig. 2e and Extended Data Fig. 2-2a-b). Remarkably, by comparing the differentiation profile of GPC4sh hiPSCs versus control, we observed a 5- to 9-time fold increase in transcript levels of all MES markers in GPC4sh hiPSCs (Fig. 2d and Extended Data Fig. 2-2a-b). Consistently, the expression of DE and ME markers was also increased of 5- to 12-folds and 5- to 19-folds, respectively, in differentiating GPC4sh cells compared to controls, (Fig. 2e and Extended Data Fig. 2-2a-b). Thus, these results indicate that GPC4sh hiPSCs acquire enhanced differentiation potential towards MES and its derivatives lineages, DE and ME, then control hiPSCs. Consistent with RT-qPCR results, immunocytochemistry revealed a ∼ 2,5 higher percentage of cells expressing the early MES marker BRACH at d1 of differentiation in GPC4sh than control cultures (Fig. 2f-g). Along the same line, the percentage of cells expressing the DE marker SOX17 and the ME marker PDGFRa at d3 of differentiation was, respectively, ∼ 17 times and ∼ 9 times higher in GPC4sh than control cultures (Fig. 2f-g). Differentiation studies performed on the second hiPSC line, AICS-0023, shown a similar increase into BRACH and SOX17 positive cells in GPC4sh AICS-0023 cells compared to controls (Extended Data Fig. 2c-d).

As DE and ME fates acquisitions are accompanied by an EMT^21, 22^, we next evaluated the capabilities of GPC4sh cells to undergo an EMT fate switch during differentiation. Marker analysis revealed that GPC4sh hiPSCs undergo a more efficient EMT transition compared to control hiPSCs, as highlighted by the increased expression of *NCAD, SNAI1, SNAI2, TWIST1* and *VIMENTIN* mesenchymal markers, and by a more efficient switch in ECAD to NCAD expression (Fig. 3a). This ECAD to NCAD transition was also marked by the concomitant appearance of SOX17 and PDGFRa proteins in NCAD positive cells (Fig. 3b-c). Altogether these results show that down-regulation of GPC4 enhances hiPSC differentiation efficiency into MES cells as well as into their DE and ME derivatives with a concomitant EMT.

**Figure 3.**
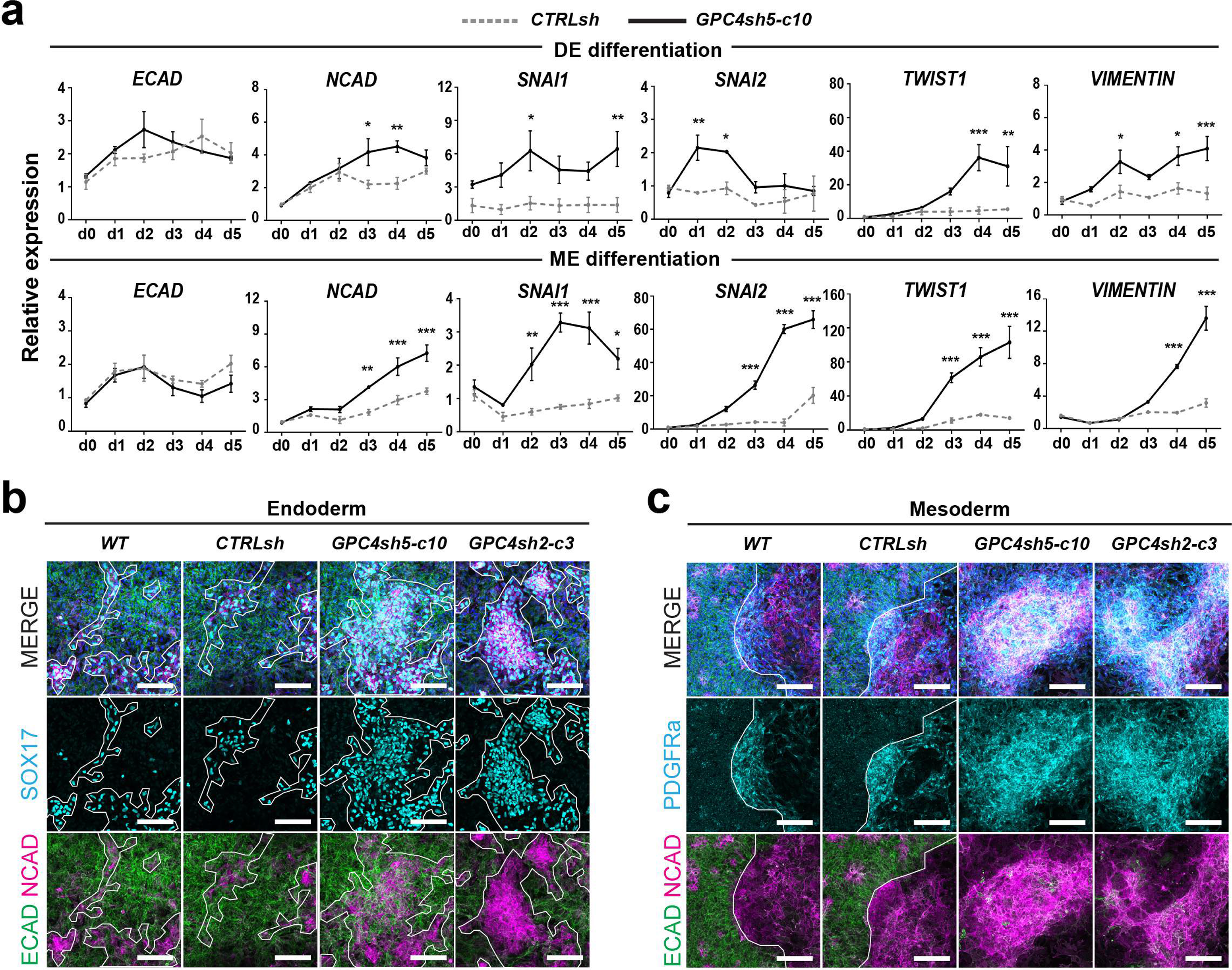
Analysis of EMT in GPC4sh cells. **(a)** RT-qPCR analysis of transcript levels of ECAD, NCAD, SNA1, SNAI2, TWIST1 and VIMENTIN in CTRLsh and GPC4sh5-c10 029 hiPSCs differentiated for 5 days into DE or ME. Top panel: DE differentiation and bottom panel: ME differentiation. Data were normalized to the d0 of CTRLsh and represented as mean ± SEM, n = 3. **(b, c)** Immunofluorescence analysis of NCAD (magenta), ECAD (green), SOX17 (cyan, DE) or PDGFRa (cyan ME) in WT, CTRLsh, GPC4sh5-c10 and GPC4sh2-c3 029 hiPSCs differentiated for 3 days into DE (b) or ME (c). Dashed areas highlight the DE and ME differentiated cells, scale bar: 100 μm.

No significant changes in the percentage of mitotic (phospho-Histone H3 positive) and apoptotic (cleaved-Caspase3 positive) cells were observed in control and GPC4sh hiPSCs (Extended Data Fig. 3a-c), thus excluding that proliferation and survival contribute to the increased differentiation efficiency in GPC4sh cultures.

### Mesendoderm fate acquisition relies on disruption of epithelial integrity

We next assessed whether disruption of the epithelial integrity in GPC4sh hiPSCs could underlie their increased differentiation potential into MES. This was addressed by restoring the epithelial integrity and polarity in GPC4sh hiPSCs, and by triggering disruption of the epithelial cell layer into CTRLsh cultures by using drugs. Drug treated cells were then exposed to ACTIVIN A to differentiate them into the MES lineage. To restore the epithelial integrity of the GPC4sh cultures, we used lyso-phosphatidic acid (LPA), a small molecule that causes an expansion of the apical domain of human neural progenitor cells^38^. Strikingly, a 24 hours treatment of GPC4sh hiPSCs with LPA was sufficient to restore a TJ organization and an epithelial integrity similar to controls (Fig. 4a). Conversely, when control hiPSCs were cultured for 24 hours in the presence of Blebbistatin (Blebbi), a drug targeting Non-Muscle Myosin II^39, 40^, the epithelial integrity, assessed by analyzing TJs, was disrupted in a fashion similar to GPC4sh cultures (Fig. 4a). Furthermore, GPC4sh cultures in which the epithelial integrity was restored by LPA lose their enhanced MES differentiation properties and behave similar to control hiPSCs, as shown by the comparable expression levels of *MIXL1, GSC* and EOMES in both cell types (Fig. 4b-c). Instead, disruption of the epithelial integrity in control hiPSCs by Blebbi treatment enhanced their capability to undergo MES differentiation as in untreated GPC4sh hiPSCs, shown by the significant increase in *MIXL1, GSC* and EOMES expression (Fig. 4b-c). Collectively, these findings show that disruption of epithelial integrity enhances efficiency of differentiation into MES lineage.

**Figure 4.**
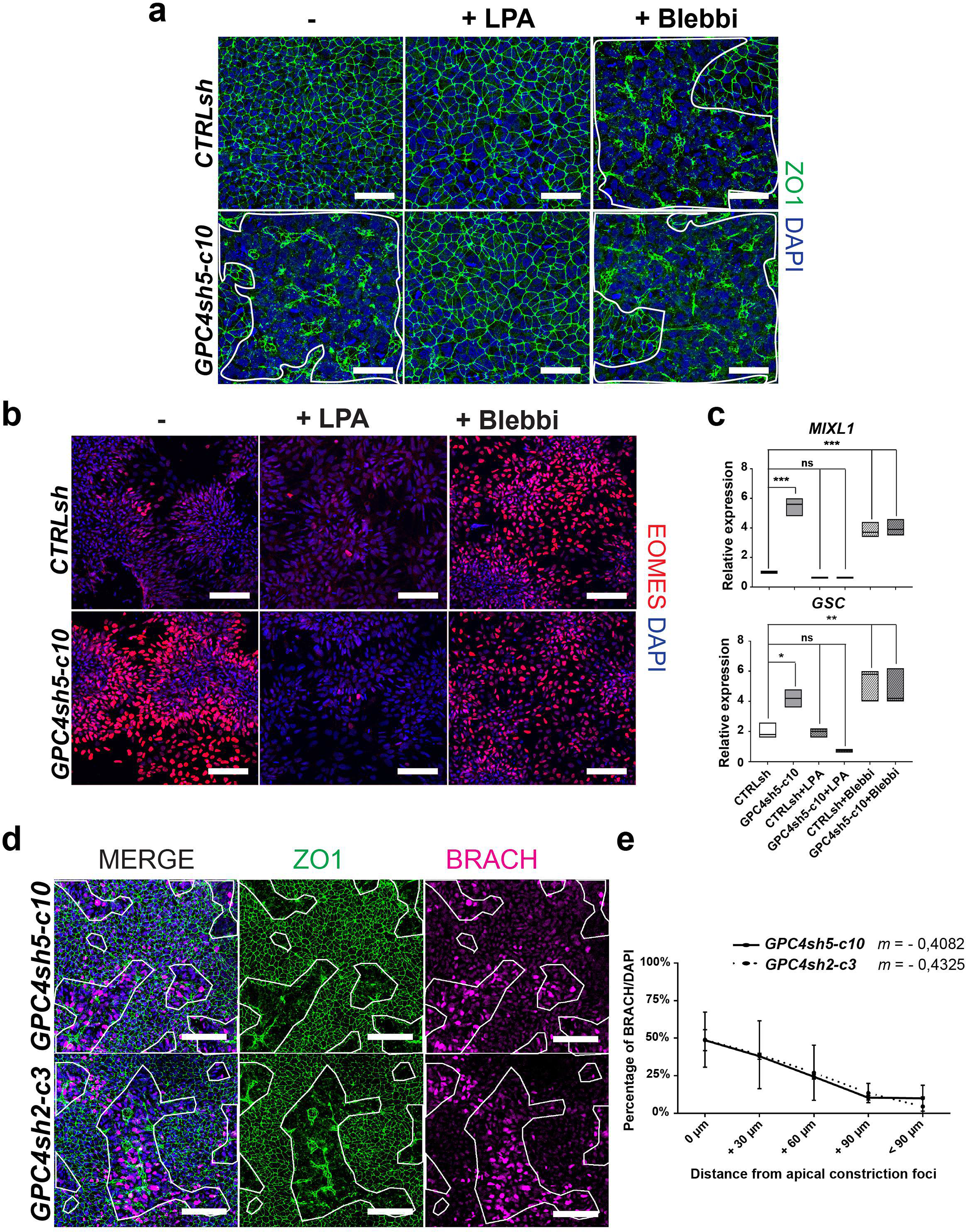
MES fate acquisition relies on disruption of epithelial integrity. **(a)** CTRLsh and GPC4sh5-c10 029 hiPSCs were treated or not with LPA or Blebbi and subjected to immunofluorescence analysis of ZO1 (green). Scale bar: 100 μm. **(b, c)** CTRLsh and GPC4sh5-c10 029 hiPSCs treated or not with LPA or Blebbi were differentiated into MES and the expression levels of MES specific markers were analyzed by (b) immunofluorescence analysis of EOMES (red; Scale bar: 100 μm.) and (c) RT-qPCR analyses of *MIXL1* and *GSC* transcript levels. Transcript levels were normalized to the mean of CTRLsh ΔCt. Box plots represent the median with min and max values, n= 3. **(d)** GPC4sh5-c10 and GPC4sh2-c3 029 hiPSCs were differentiated into MES with ACTIVIN A and subjected to immunofluorescence analysis of ZO1 (green) and BRACH (magenta). Dashed areas highlight the extent of zones with disrupted TJs. Scale bar: 100 μm. **(e)** Percentage of BRACH positive cells as a function of distance from the zone with disrupted TJs were quantified from staining shown in (d). 0 μm indicates cells located within the zones of disrupted TJs, + 30 μm, + 60 μm, + 90 μm and < + 90 μm indicates cells located at 0 to 30µm, 30 to 60µm, 60 to 90µm or more than 90µm away from zones of disrupted TJs, respectively. Slope of the curves are represented as m. Data are represented as mean ± SD. n = 3. Statistical analysis for the overall figure: (c) one-way ANOVA followed by Sidak’s multiple comparison test, (d) linear regression analysis. P values: (***) < 0.001, (**) < 0.01, (*) < 0.05.

Interestingly, by comparing the spatial distribution of cell patches with altered TJ organization with that of BRACH expressing cells in differentiating GPC4sh cultures at MES stage, we found a striking correlation into their spatial distributions. The correlation coefficient, calculated by quantifying the percentage of BRACH positive cells as a function of distance from epithelial disrupted areas (Fig. 4d-e), revealed a positive relationship between the two variables with a slope ranging from −0,4082 to - 0,4325 for GPC4sh5 and GPC4sh2 lines, respectively. (Fig. 4d-e). Taken together these results show that local disruption of epithelial integrity in the hiPSC layer regulates the onset of MES fate acquisition and its spatial pattern.

### Disruption of hiPSC epithelial integrity enables activation of BMP and ACTIVIN/NODAL signaling

Decades of research on mammalian gastrulation have revealed that MES formation and spatial patterning is under the control of a signaling cascade initiated by BMP and involving activation of WNT, and NODAL pathways^41–43^. Our results show that disruption of epithelial integrity in GPC4sh hiPSCs does not prime hiPSCs for MES fate acquisition. Yet, GPC4sh hiPSCs undergo efficient differentiation towards the MES lineage following ACTIVIN A stimulation. This raised the question of whether these structural changes in hiPSC epithelial layer may enhance their ability to activate BMP, ACTIVIN/NODAL and/or WNT-βcat signaling cascade. In agreement with this possibility, prior studies using hESCs underscored as the epithelial organization and polarity of ESCs prevents accessibility of TGF-β superfamily ligands to their receptors located at the basolateral cell membrane as well as induction of the WNT-βcat signaling pathway^41–43^. As above reported, disruption of epithelial integrity in GPC4sh hiPSCs alters the apico-basal polarity of epithelial cells thus enabling accessibility to proteins otherwise localized at the basolateral cell side (Fig. 1g and Extended Data Fig. 1f).

To answer to this question, we analyzed the capacity of control and GPC4sh hiPSCs grown in self-renewal conditions to activate the BMP and ACTIVIN pathways upon 1-hour stimulation (immediate response). Cells responding to BMP4 and ACTIVIN A stimulations were visualized by immunocytochemistry of nuclear phospho-SMAD1/5 (pSMAD1,5 S463/S465) and phospho-SMAD2 (pSMAD2 S465/S467), respectively^30^. After 1-hour of stimulation, we found that the percentages of cells activating BMP and ACTIVIN signaling were ∼ 3 and ∼ 2 folds higher in GPC4sh compared to control hiPSCs, respectively (Fig. 5a-b and Extended Data Fig. 5a-b). Additionally, image analysis revealed also that most cells responding to BMP4 and ACTIVIN A stimulations localized in the areas with disrupted TJs (Fig. 5a and Extended Data Fig. 5a). This was confirmed by quantifying the percentage of pSMAD1,5 or pSMAD2 positive cells and their distance from epithelial disrupted areas. Strikingly, this quantification analysis revealed a good positive correlation between these two variables, showing that they are interconnected (Fig. 5c and Extended Data Fig. 5c). Analysis of the second hiPSC line, AICS-0023, confirmed that most cells responding to BMP stimulation were localized in the areas with disrupted TJs in GPC4sh cultures compared to control cells (Extended Data Fig. 5d).

**Figure 5.**
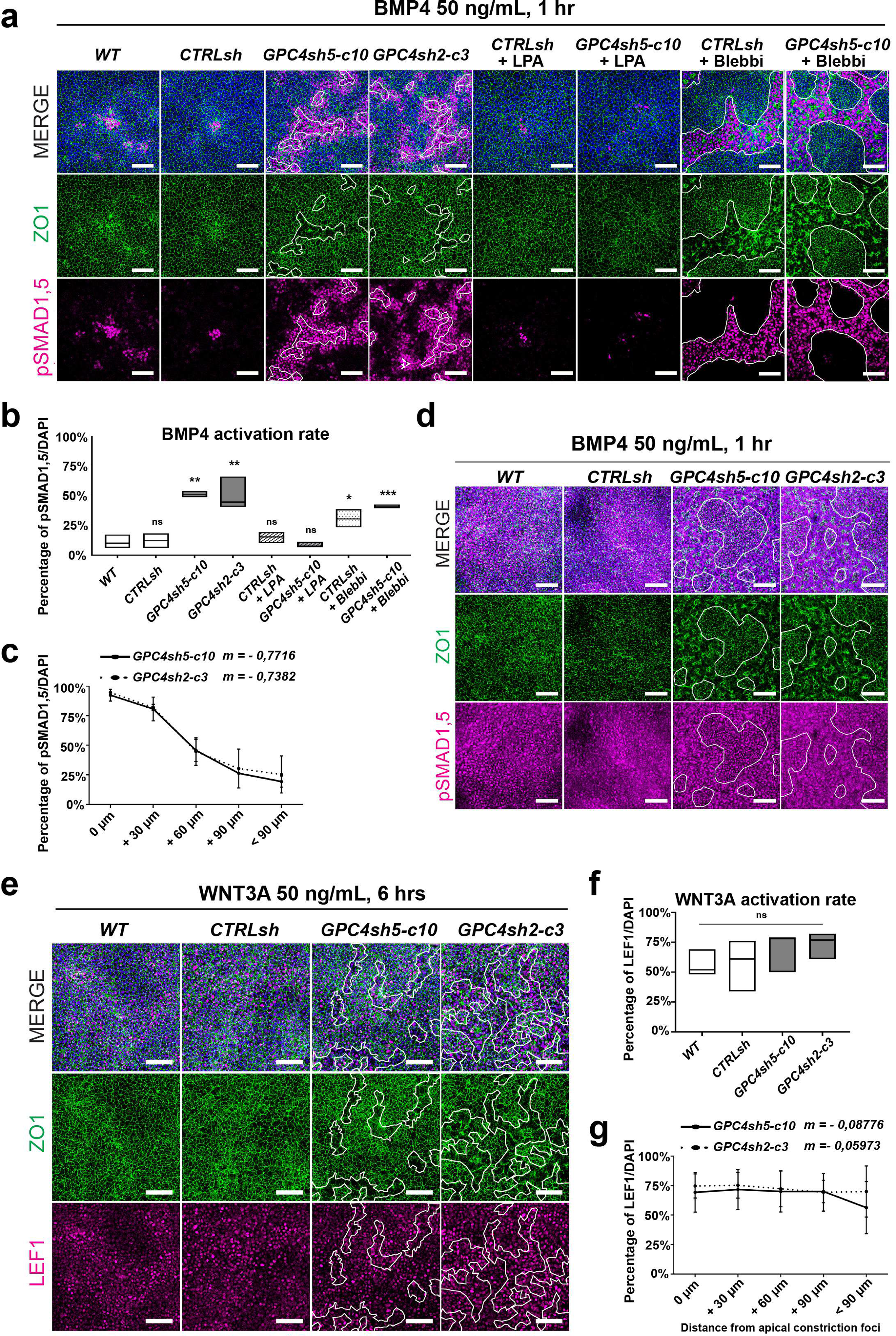
Disruption of epithelial integrity enables activation of TGF-β signaling. **(a)** Cultures of WT, CTRLsh, GPC4sh5-c10 and GPC4sh2-c3 029 hiPSCs treated or not with LPA or Blebbi were stimulated for 1 hour with 50 μg/mL of BMP4, and analyzed for pSMAD1,5 (magenta) and ZO1 (green) proteins. Dashed areas highlight zones with disrupted TJs, scale bar: 100 μm. **(b)** Percentages of pSMAD1,5 positive cells were quantified from staining shown in (a). Box plots represent the median with min and max values, n = 3. **(c)** Percentages of pSMAD1,5 positive cells as a function of distance from zones with disrupted TJs quantified in GPC4sh5-c10 and GPC4sh2-c3 029 hiPSCs from staining in (a). 0 μm indicates cells located within zones with disrupted TJs, + 30 μm, + 60 μm, + 90 μm and < + 90 μm indicates cells located at 0 to 30µm, 30 to 60µm, 60 to 90µm or more than 90µm away from the zone with disrupted TJs, respectively. Data are represented as mean ± SD, n = 3. **(d)** Immunofluorescence analysis of pSMAD1,5 (magenta) and ZO1 (green) in WT, CTRLsh, GPC4sh5-c10, GPC4sh2-c3 029 hiPSCs grown in transwell plates and stimulated from the basal side for 1 hour with 50 μg/mL of BMP4. Dashed areas highlight the extent of zones of disrupted TJs, scale bar: 100 μm. **(e)** Cultures of WT, CTRLsh, GPC4sh5-c10 and GPC4sh2-c3 029 hiPSCs were stimulated for 6 hours with 50 μg/mL of WNT3A, and analyzed for LEF1 (magenta) and ZO1 (green) proteins. Dashed areas highlight the disturbed TJs, scale bar: 100 μm**. (f)** Percentages of LEF1 expressing cells were quantified from staining in (d). Box plots represent the median with min and max values, n = 3 **(j)** Percentages of LEF1 expressing cells as a function of the distance from zones with disrupted TJs quantified in GPC4sh5-c10 and GPC4sh2-c3 029 hiPSCs from staining shown in (d). 0 μm indicates cells located within the zones of TJs disruption, + 30 μm, + 60 μm, + 90 μm and < + 90 μm indicates cells located at 0 to 30µm, 30 to 60µm, 60 to 90µm or more than 90µm from the zone of TJs disruption respectively. Data are represented as mean ± SD, n = 3. Statistical analysis for the overall figure: (c, f) linear regression analysis, (b, e) one-way ANOVA followed by Dunnett’s multiple comparison test. P values: (***) < 0.001, (**) < 0.01, (*) < 0.05.

In line with these results, we found that when epithelial integrity of GPC4sh cultures was restored by LPA, the percentage of nuclear pSMAD1,5 positive cells was reduced to that of control hiPSCs (Fig. 5a-b). Conversely, when the epithelial integrity of control cells was disrupted by Blebbi treatment, the percentage of cells with activated BMP signaling was similar to that of GPC4sh5 hiPSCs and they mainly localized to areas with disrupted TJs (Fig. 5a-b). Finally, we observed a global activation of the BMP signaling in control and GPC4sh hiPSCs, when BMP4 was delivered from the basal side (Fig. 5d). These findings suggest that epithelial integrity in control hiPSCs prevents BMP4 signaling activation when ligands are provided from the apical side, likely due to inaccessible receptors. Taken together, these results show that local disruption of epithelial integrity enables perception of BMP4 and ACTIVIN A signaling proteins by cells located in these area and activation of TGF-β signaling in a spatially regulated manner. This event will in turn trigger expression of MES markers in cells as well as their spatial pattern (Fig. 4b-e).

Considering that WNT-βcat signaling is an additional crucial regulator of MES differentiation, we assessed whether epithelial integrity influence the hiPSC ability to activate WNT-βcat pathway upon stimulation. For these studies, we used immunocytochemistry to analyze the percentage of cells expressing LEF1, a co-factor of β-cat that is a direct target of WNT signaling^44, 45^. Control and GPC4sh hiPSCs were stimulated for 6 hours with the WNT3A ligand. By following LEF1 distribution, we observed ∼ 60% of cells in both control and GPC4sh hiPSCs expressing LEF1, detected in a salt and pepper manner (Fig. 5e-f). Finally, there was no correlation between localization of LEF1 positive cells and patches of disrupted epithelial integrity in GPC4sh cultures (*m* _GPC4sh5_ = −0,08776 and *m* _GPC4sh2_= −0,05973) (Fig. 5g). Thus, these results indicate that epithelial integrity does not impact on activation of WNT-βcat signaling nor its spatial localization.

### Disruption of epithelial integrity prolongs TGF-β signaling activation

Recent studies using micropattern devices showed that the cellular response of PSC colonies to BMP stimulation varies over time. Indeed, a gradual restriction of BMP4 responding cells in micropattern going from all over the colony to the edge over time was found, as cell density increases, which will then prolong the induction time in the cells at the edge^11^. We therefore asked whether disruption of epithelial integrity influences maintenance of TGF-β signaling over time in hiPSC GPC4sh versus control cultures. Cells were plated at the density used for differentiation experiments (1,1×10^5^ cells/cm²) and stimulated for 24 hours with BMP4. The kinetic of BMP4 pathway response was then followed at 2, 6, 12, 18 and 24 hours by using pSMAD1,5 immunocytochemistry. After 2 hours of stimulation, when hiPSC cultures do not yet form a dense epithelial sheet, we observed a similar BMP4 response in both CTRLsh and GPC4sh5 hiPSCs with ∼ 80% of pSMAD1,5 cells positive. However, as time progresses and cultures become denser, we found that the percentage of cells with BMP activation decreased rapidly in CTRLsh cultures compared to GPC4sh with solely ∼ 5% of pSMAD1,5 positive cells after 12 hours of stimulation (Fig. 6a-b). This lower percentage of pSMAD1,5 positive cells in CTRLsh cultures persisted at later time points (Fig. 6a-b). Similar effects were observed when CTRLsh and GPC4sh5 hiPSCs were stimulated with ACTIVIN A (Extended Data Fig. 6a-b). Strikingly, loss of epithelial integrity correlated precisely with the spatial activation of BMP4 signaling in long-term GPC4sh cultures (Fig. 6c). These results show that disruption of epithelial integrity enables a prolonged activation of TGF-β signaling in cultured cells, which might in turn impact on robustness of cell fate acquisition. Outcomes provide also new mechanistic insights on how MES triggering signals could be temporally controlled.

**Figure 6.**
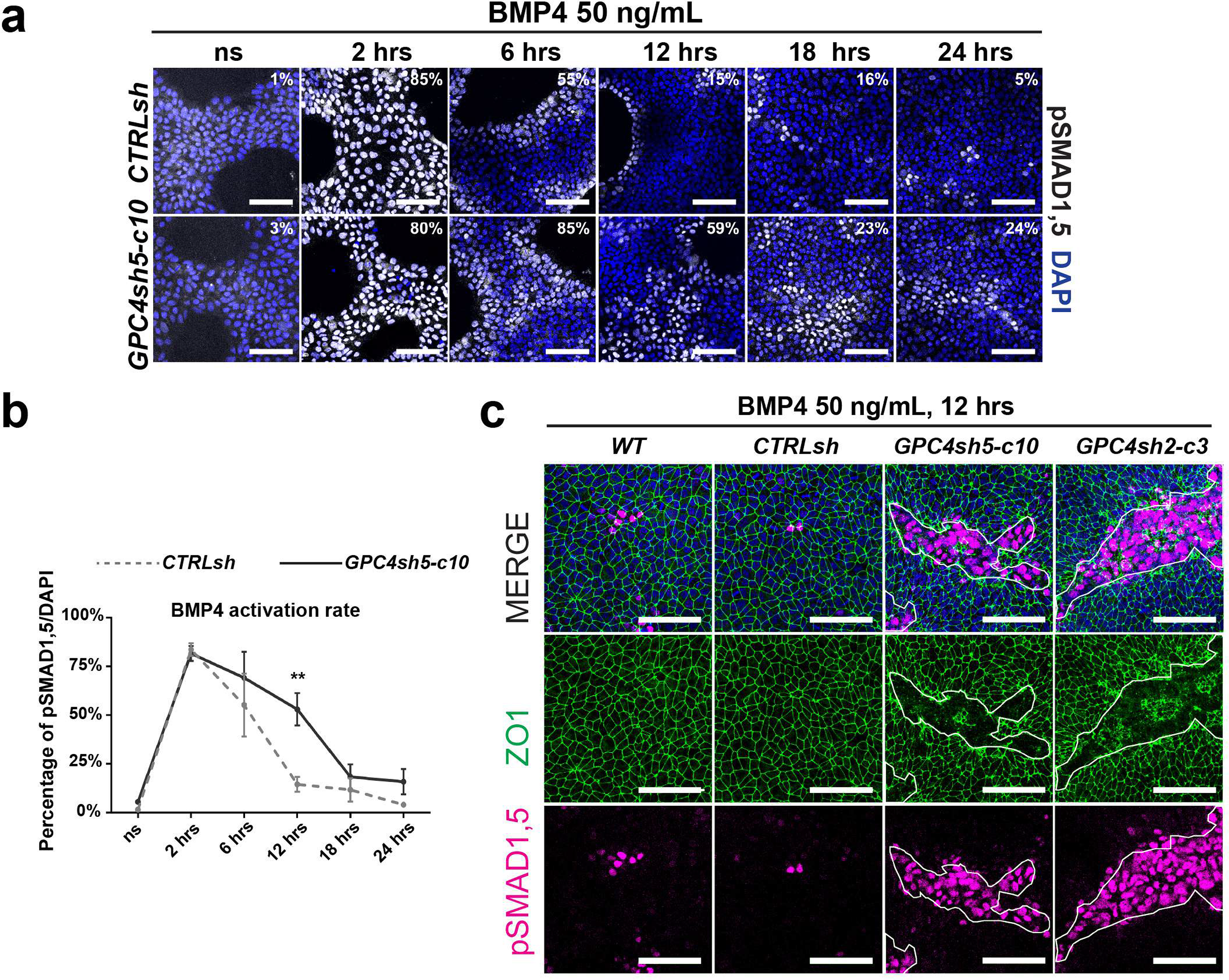
Disruption of epithelial integrity prolongs TGF-β signaling activation. **(a)** Cultures of CTRLsh and GPC4sh5-c10 029 hiPSCs were stimulated for 2, 6, 12, 18 and 24 hours with 50 μg/mL of BMP4, and subsequently subjected to immunofluorescence analysis of pSMAD1,5 (grey). Scale bar: 100 μm. **(b)** Percentages of pSMAD1,5 expressing cells were quantified from staining shown in (a). Data are represented as mean ± SEM, n = 3. **(c)** Cultures of WT, CTRLsh, GPC4sh5-c10 and GPC4sh2-c3 029 hiPSCs were stimulated for 12 hours with 50 μg/mL of BMP4, and subsequently subjected to immunofluorescence analysis of ZO1 (green) and pSMAD1,5 (magenta). Dashed areas highlight the extent of zones of disrupted TJs, scale bar: 100 μm. Note the correlation of disrupted epithelial area and the presence of pSMAD1,5 positive cells in GPC4sh cultures. Statistical analysis for the overall figure: (b) two-way ANOVA followed by Sidak’s multiple comparison test. P values: (***) < 0.001, (**) < 0.01, (*) < 0.05.

## DISCUSSION

Unravelling the processes of primitive streak formation and epiblast patterning in mammals remains a great challenge, especially for human embryos. In vitro models of the human epiblast cells based on hPSCs are offering the unique opportunity to decompose these complex processes and allow independent manipulation of the underlying mechanisms. Current consensus focuses on the concept that these early embryonic processes are controlled by feedback interactions between biochemical signals (morphogen and their inhibitors/activators) and tissue organization (epithelial cell polarity, adhesion and cytoskeleton tension). In the present studies we explored how these instructive cues functionally interact and whether they operate into a hierarchical manner. Here we report that hiPSCs with reduced protein levels of the morphogen regulator GPC4 represent a unique cellular system to investigate the physical effects of disrupted epithelial cell polarity and architecture on cellular response to TGF-β proteins at the human MES onset. We demonstrated that GPC4 down-regulation in hiPSCs results into a distinct morphological organization of hiPSC layer wherein alteration of TJs leads to a mosaic hiPSC structural pattern with areas of disrupted epithelial integrity. This phenotype alters the apico-basal cell axis causing exposure of proteins in the basolateral cell compartment. Moreover, it influences the spatial regulation of cell fate acquisition, as cells within areas of disrupted epithelia integrity become sensitive to BMP4 and ACTIVIN A protein stimulation and acquire a MES fate (e.g., BRACH and EOMES expression). Thus, our results demonstrate as distinct changes in epiblast cell polarity and tissue integrity can foster the spatial response of epiblast cells to the MES-initiating signal and the subsequent differentiation onset into a precisely controlled spatial pattern.

Our experiments involving apical and basal stimulation of CTRLsh and GPC4sh hiPSCs with BMP4 and ACTIVIN A proteins provide insights into the underlying mechanisms. Under culture conditions designed for pluripotency maintenance, TGF-β reception in CTRLsh hiPSCs is restricted to the basolateral cell domain, as cells activate TGF-β signaling only following basal ligand stimulation. This is consistent with recent reports in which the use of 2D micropatterned hESC-based epiblast model revealed that hESCs localize their BMP receptors to the basolateral domain just below the tight junctions, making them non-responsive to apically applied BMP4^11^. In contrast, the disrupted epithelial integrity of GPC4sh hiPSCs enables local perception of BMP4/ACTIVIN A and signaling pathway activation independently of whether the ligand is supplied from the basal or apical cell side. Thus, the morphological organization of the GPC4sh hiPSC layer with disrupted TJs enables increase accessibility of ligand to receptors. A similar pattern of basolateral BMP4 receptor localization below TJs has also been reported in vivo in the developing mouse epiblast^6^. Similar to the finding we describe here, lack of TJs at the border between the extraembryonic ectoderm and posterior epiblast enables BMP ligands secreted into the proamniotic cavity to access receptors at the posterior epiblast edge, and to trigger PS formation at this level^6^. Our results together with these embryological studies point to a feedback loop between epiblast morphology and signaling, in which epiblast epithelial integrity controls spatial activation of TGF-β signaling and patterning. In this context, GPC4sh hiPSCs may represent an optimal human cellular system to analyse further this feedback interaction and to experimentally dissect mechanisms controlling epiblast cell polarity and tissue morphology. As disruption of epithelial integrity in GPC4sh hiPSCs extend the duration of BMP and ACTIVIN signaling pathway activation, the GPC4sh hiPSCs could also enable a study of the potential role of epithelial morphology/organization into the temporal regulation of MES initiating signals.

In contrast to TGF-β proteins, disruption of epithelial integrity does not regulate spatial activation of WNT-βcat signaling in our experimental conditions, as shown by a rather homogeneous activation of LEF1 within the GPC4sh hiPSC layer following WNT3A stimulation. These results are consistent with previous works showing that WNT receptors are accessible to WNT ligands from both the apical and the basolateral cell side ^46^. Nevertheless, activation of WNT-βcat signaling is sensitive to changes in tissue-level forces, which are dependent on tissue organization. As modification of TJ distribution affects global tissue tension, we cannot exclude that with our experimental conditions prevented detection of altered hiPSC response to WNT3A stimulation associated with of epithelial integrity. Our studies were conducted on cells cultured on stiff substrates such as glass or plastic surfaces on which possible changes in tissue tension induced by loss of TJs might be buffered. Studies by others have shown that in hESCs cultured on soft substrates, activation of Wnt-βcat signaling correlates with high tissue-level forces^13, 47^. Thus, it could be relevant to perform future studies culturing GPC4sh hiPSCs on soft substrates, such as hydrogels surfaces, to assess whether disruption of epithelial integrity modulate WNT response by modifying global tissue tension.

Interestingly, disruption of epithelial integrity in the GPC4sh hiPSC layer does not perturb self-renewal and pluripotency neither promotes premature expression of EMT markers or lineage specific genes (present studies and ref. 14). Thus, alterations in TJs, although priming cells towards morphogen perception, are not sufficient to prime differentiation or lineage fate choices. It is tempting to speculate that this epithelial disruption is not sufficient to achieve a threshold level required for differentiation. Alternatively, unknown factor(s) could buffer such drastic alteration in epithelial organisation. The observed maintenance of stemness/pluripotency in GPC4sh hiPSCs is consistent with recent studies on hESCs carrying a genetic knockout of the human cell-cell adhesion protein, E-CAD. Nevertheless, these cells show a transient activation of lineage specific genes and preferential differentiation towards MES, thus suggesting that loss of E-CAD impact on lineage fate decisions^47, 48^.

Our results disclose also a new role of GPC4 in maintaining epithelial cell integrity. Previous studies performed on Zebrafish and Xenopus embryos have highlighted GPC4 function during ME convergence extension movements at gastrulation in which GPC4 controls polarity and directed migration of mesodermal cells^18, 19, 49, 50^. Moreover, in Zebrafish embryos GPC4 promotes DE cell polarity by influencing localization of N-CAD on the cell surface through regulation of Rab5c-mediated endocytosis^51^. Future experimental settings will clarify whether mechanisms involving GPC4-mediated endocytosis of membrane proteins participating to cell-cell and cell-matrix contacts underlie the epithelial integrity defects found of GPC4sh hiPSCs.

In conclusion, our results highlight a new mechanism by which tissue architecture can restrict morphogen sensing and position cell fate changes within space.

Our differentiation analysis of GPC4sh hiPSCs reveals their increased differentiation propensity in ME and DE compared to conventional hiPSCs. These finding support that possibility that targeting GPC4 in different hiPSCs may provide a new strategy to overcome the technical difficulties involved in their differentiation, thus providing a tool to promote hiPSC application for disease modelling, drug screening and regenerative medicine. Yet, the GPC4sh hiPSCs may provide a platform where differentiation relevant processes can be followed and deconstructed. GPC4sh hiPSCs are thus a mean to get new insights into developmental processes and for advancing into regenerative medicine. We expect that this mechanism of modulating epithelial integrity by altering TJs will be of more general relevance in other physiological and pathological events in which there is a high degree of epithelial cell plasticity such as remodeling of the epithelial barriers, tissue inflammation and tumorigenesis.

## METHODS

### Human induced pluripotent stem cell lines and culture

WT 029 hiPSCs were a courtesy of Michael Kyba (University of Minnesota)^14, 52^. The AICS-0023 previously used were from Coriell Institute for Medical Research. The generation of GFPsh hiPSCs and GPC4sh hiPSCs were generated as previously described (Corti et al 2021). hiPSCs were maintained on Matrigel-coated plates (Corning, BV 354277) within mTeSR1 medium (mTeSR™1 (Stemcell Technologies 85850), 1% Penicillin/Streptomycin (Invitrogen, ref. 15140122). In order to maintain hiPSC cultures not exceeding 80% of confluency, cells were passaged at 1/8 ratio every 3 or 4 days with Accumax (Millipore, ref. SCR006).

### Alkaline phosphatase assay

Alkaline Phosphatase (AP) detection has been performed with the Alkaline Phosphatase Staining Kit II (Stemgent, ref. 00-0055). Briefly, hiPSCs were seeded on Matrigel-coated coverslips until reaching a maximum of 80% of confluency. Cells were incubated with Fix Solution at room temperature for 5 minutes and then washed with PBS. Afterwards, Cells were incubated with AP Substrate solution in the dark, at room temperature for 10 minutes. The reaction was stopped by aspirating the AP Substrate solution and washing the wells twice with PBS. Images were captured on a Zeiss Stereo microscope.

### Cell cycle analysis

hiPSCs were seeded on Matrigel-coated plates and analyzed at 30-50% of cells confluency. Nuclei were extracted with lysis buffer (Chemometec) and incubated with 10 µg/ml of DAPI for 5 minutes at 37°C. Nuclear DAPI intensity was measured with the Cytometer NucleoCounter® NC-3000™ (Chemometec) and cell cycle was analyzed with the NucleoView NC-3000™ program (Chemometec).

### Differentiation assays into MES, DE and ME lineages

Differentiation experiments of hiPSCs into DE and ME lineages were performed by following the protocols published by D’amour et al., 2005 and Laflamme et al., 2007 respectively. To generate MES, DE and ME lineages, hiPSCs were seeded on Matrigel-coated plates or coverslips at a density of 1.1×10^5^ cells/cm² in mTeSR1 medium supplemented with 10 μM of ROCK inhibitor Y-27632 (Ri; Tocris, ref. 1254). To induce MES differentiation, hiPSCs were washed twice with RPMI (Invitrogen, ref. 21875034) to remove self-renewal growth factors and treated for 1 day with ACTIVIN A (R&D, ref. 338-AC, 100 ng/mL) in DE medium (RPMI 1640, 1% Penicillin/Streptomycin, 1% L-Glutamine (Gibco, ref. 25-030-081). DE differentiation was induced by threating MES differentiated cultures for 2 days with ACTIVIN A (R&D, ref. 338-AC, 100 ng/mL) in DE medium supplemented with 0,2% of FBS (Hyclone, ref. SH30071.02E). ME differentiation was induced by threating PS differentiated cultures for 2 days with BMP4 (R&D, ref. 314-BP, 10 ng/mL) in ME differentiation medium (RPMI 1640, 2% B27 without Insulin (Gibco, ref. A1895601), 1% Penicillin/Streptomycin, 1% L-Glutamine (Invitrogen, ref. 25030149). Medium was replaced every day. The protocol reporting medium replacement is summarized in Supplementary Table 3.

For experiments involving drug treatments, hiPSCs were seeded on Matrigel-coated plates or coverslips at a density of 0,25×10^5^ cells/cm² within mTeSR1 medium supplemented with Ri. Rescue of epithelial integrity was induced by threating hiPSCs for 1 day with LPA (StemCell Technologies, ref. 72694, 5 µM). Chemical disruption of epithelial integrity was induced by threating hiPSCs for 1 day with Blebbistatin (Sigma, ref. B0560, 10 µM). MES differentiation was subsequently induced by washing cultures twice with RPMI and threating hiPSCs for 1 day with ACTIVIN A (100 ng/mL) in DE medium supplemented with LPA (5 µM) or Blebbistatin (10 µM). Medium was replaced every day. The protocol reporting medium replacement is summarized in Supplementary Table 4.

### Stimulation assays

Stimulation assays were performed on confluent hiPSC cultures maintained in mTeSR1 medium. HiPSCs were seeded at 1.5×10^5^ cells/cm² on Matrigel-coated coverslips in mTeSR1 medium supplemented with Ri. When confluent, hiPSCs cultures were stimulated for 1 hour with BMP4 (R&D, ref. 314-BP, 50 ng/mL) or ACTIVIN A (100 ng/mL), and 6 hours with WNT3A (R&D, ref. 5036-WNT, 50ng/mL) in mTeSR1 medium. Cells were then fixed and signaling pathways activation was assessed trough immuno-cytochemical analyses. Medium was replaced every day. Medium replacement protocols are summarized in Supplementary Table 5.

For experiments of rescue and chemical induction of epithelial integrity, hiPSCs were seeded on Matrigel-coated coverslips at a density of 0,75×10^5^ cells/cm² in mTeSR1 medium supplemented with Ri. The next day, medium was replaced by fresh mTeSR1 medium. The day after, medium was replaced for 1 day by mTeSR1 medium supplemented with 5 µM of LPA or 10 µM of Blebbistatin. Finally, confluent cultures were stimulated for 1 hour with BMP4 (50 ng/mL) in mTeSR1 medium, and then fixed and analyzed by immunocytochemistry.

For long-term stimulation experiments of BMP and ACTIVIN pathways, hiPSCs were seeded on Matrigel-coated coverslips at a density of 1,1×10^5^ cells/cm² in mTeSR1 medium supplemented with Ri. The next day, medium was replaced by fresh mTeSR1 medium. The day after, hiPSCs cultures were stimulated with BMP4 (50 ng/mL) or ACTIVIN A (100 ng/mL) in mTeSR1 medium. hiPSCs cultures were then fixed and analyzed by immunocytochemistry at 0, 2-, 6-, 12-, 18- and 24-hours following stimulation.

### RNA isolation, cDNA synthesis and quantitative PCR

Total RNA was isolated with RNeasy Min Elute clean-up kit (Qiagen, ref. 74204), according to the manufacturer’s protocol. Concentration and RNA quality were evaluated by NanoDrop (ND-1000 Spectrophotometer; Thermofisher Scientific). 600ng of RNA was reverse-transcribed (RT) to generate cDNA using iScript reverse transcription kit (Biorad, ref. 170-8841), following manufacturer’s instruction. 2,7 ng of cDNA were amplified by quantitative PCR (qPCR) using SYBR Green qPCR SuperMix-UDG with Rox (Thermofisher, ref. 11733046) and 0.1µM of forward and reverse primers. Levels of transcripts (Ct) were normalized to those of the housekeeping gene GAPDH (ΔCt) and subsequently to the mean ΔCt of the reference sample (ΔΔCt). Results were reported either as relative quantities (RQ = 2^-ΔΔCt) or as fold change (FC = -ΔΔCt). The sequence of forward and reverse primers used for qPCR analyses are listed in Supplementary Table 6.

### RNA sequencing analyses

For RNA-seq, 4×10^4^ cell/cm^2^ cells of WT, shCTRL and shGPC4-1 029 hiPSCs were seeded in triplicate in 12-well plates. After three days they were dissociated for RNA extraction as described above. RNA quality was then tested with the instrument 2100 Bioanalyzer Agilent (Purity: OD 260/280: 1.8 - 2.0; OD 260/230: 2.0 - 2.2; RIN or RQI value ≥ 8). 1 μg of RNA per sample at the Concentration: > 20 ng/μl dissolved in RNase-, DNase- and protease-free molecular grade water was sent to GATC Biotech. The RNA-sequencing was performed by using Illumina HiSeq 2500 technology. The quality of the raw reads was assessed using FastQC v0.11.9 toolkit (Andrews, 2010). Adapters and low-quality reads were trimmed using Trimmomatic v0.39 (Bolger et al., 2014). Paired-end reads were aligned with HiSat2 v2.2.1 (Kim et al., 2015) using default options. Gene counts were quantified using feature Counts from the Subread v2.0.1 package (Liao et al., 2019). Alignment and gene counts were generated against the GRCh38.p13 (Ensembl release 101). The low expressed genes which did not have more than one count per million reads (1CPM) in at least three samples within each dataset were removed from further analysis. Gene counts were then normalized and used for differential expression testing using DESeq2 v1.28.0^53^ (Supplementary Tables 1,2).

### Western Blots

For western blots, cells were lysed in 1X EBM lysis buffer in the presence of a protease and phosphatase inhibitors mixture. 20 to 50 μg of whole cell lysates were resolved on 10% Anderson gels and transferred to PVDF membranes. After blocking with 5% milk in PBS, 0.1% Triton X100, the membranes were incubated overnight at 4°C with primary antibodies. Primary antibodies used are listed in Supplementary Table 7. The membranes were then washed, incubated with anti-rabbit or anti-mouse IgG-peroxidase (1:4000, Jackson, ref. 211-035-109) at room temperature for 1hr, and peroxidase activity was visualized with the ECL Plus Kit (Amersham).

### Immuno-cytochemical analyses

Cells were fixed in 4% paraformaldehyde (PFA) for 10 minutes and washed three times with PBS. Permeabilization was done by incubating the cells with 0.3% or 0,1% Triton X100 in PBS for 20 minutes. Non-specific interactions were blocked for 1 hour with blocking solution (3% BSA, Sigma ref. A9647, 2% donkey serum, Abcam ref. ab7475, 0,3% Triton X-100, PBS). Primary antibodies were diluted in the blocking solution and incubated overnight at 4°C. Primary antibodies used for immuno-cytochemical analyses are listed in Supplementary Table 8. Secondary antibodies were diluted at 1/500 in blocking solution and incubated for 1 hour at room temperature. Nuclei were stained with DAPI (1/1000 in PBS) for 5 minutes. Finally, coverslips were mounted on slides using Fluoromount-G mounting medium. Picture were acquired with a confocal Zeiss LSM 880 microscope and analyzed through FIDJI, ZEN and AMIRA software.

### Quantifications of immunostaining

#### Signal quantification and count

For differentiation and stimulation assays, number of positive cells showing nuclear localization of transcription factors (LEF1, BRACH and SOX17), downstream effectors (pSMAD2 or pSMAD1), were automatically counted by FIDJI. Shortly Median filtering, automatic Huang thresholding and watershed treatment were used to generate a MASK of Nuclei (DAPI) and to count their numbers. For the nuclear marker of interest, background signal was reduced by a Rolling Ball method. Then, Top hat filtering and automatic Otsu thresholding were used to generate a MASK of the nuclear marker and to count positive cells. For cytoplasmic signal (PDGFRa), the number of positive were manually counted on FIDJI.

#### Morphological analysis

To quantify disrupted epithelial areas, ZO1 immunostaining was manually analyzed with FIDJI, and a MASK corresponding to the disrupted area was generated. From this MASK, 4 others concentric MASKs corresponding to 4 localizations, corresponding to 0 to 30µm, 30 to 60µm, 60 to 90µm or more than 90µm away from disrupted areas, were generated and used to analyze the correlation between cells positive for pSMAD1,5, pSMAD2, LEF1 or BRACH (as described in signal quantification and count) and their position relative to the disrupted areas.

### Statistical analyses

All statistical analyses were performed with GraphPad Prism version 8 software. All statistical test used are indicated in the respective figure legends and data were presented as median ± standard error of the mean (SEM), unless stated otherwise. Statistics were reported as: ns=not significant, * = pvalue < 0.05, ** = pvalue < 0.01, *** = pvalue < 0.001.

### Data availability

The RNA-seq data are available on Zenodo (https://zenodo.org/) with the DOI 10.5281/zenodo.5517383.

### Code availability

The workflow allowing to achieve the differential expression analysis of RNA-seq data is available on Zenodo (https://zenodo.org/) with the DOI 10.5281/zenodo.5517512 and is under the BSD-3 license. For the differential RNA-seq analyses, the genes with a log2fc between −0.58 to 0.58 and an adjusted p-value (padj) > 0.01 were not considered as significantly expressed.

## Author Contributions

D.R., T.L., and R.D. designed the research; F.M. provided input on experiment and data. D.R. and T.L. conducted experiments; D.R., T.L., and R.D. analyzed data; T.V. analyzed the RNAseq analysis; D.R. and T.L. produced initial images for figures; and D.R. and R.D. wrote the paper with assistance from all authors. D.R., T.L., T.V., F.M. and R.D. manuscript review and editing. R.D. provided funding.

## ACKNOWLEDGEMENTS

We thank: all members of our labs for helpful discussions and comments; Robert Kelly, Jack Llewellyn and Benoit Sorre for critical reading of the manuscript; S. Corti for contribution during the preparation of RNA for RNAseq studies, E. Bazellières for discussions and reagents. Microscopy was performed at the imaging platform of the IBDM, supported by the ANR through the “Investments for the Future” program (France-BioImaging, ANR-10-INSB-04-01). The authors thank members of the IBDM core facility for microscopy and mouse husbandry for their technical support. This project is a part of the research program of the Centre of Excellence DHUNE, which is supported by the French National Plan on Neurodegenerative Diseases funded by the French Ministry of national education, higher education and research and the «Investissements d’Avenir» French Government program. The IBDM is affiliated with NeuroMarseille, the AMU neuroscience network, and with NeuroSchool, the AMU graduate school in neuroscience supported by the A*MIDEX foundation (AMX-19-IET-004) and the “Investissements d’Avenir” program (nEURo*AMU, ANR-17-EURE-0029 grant).

## EXTENDED DATA

**Extended Data Fig. 1.**
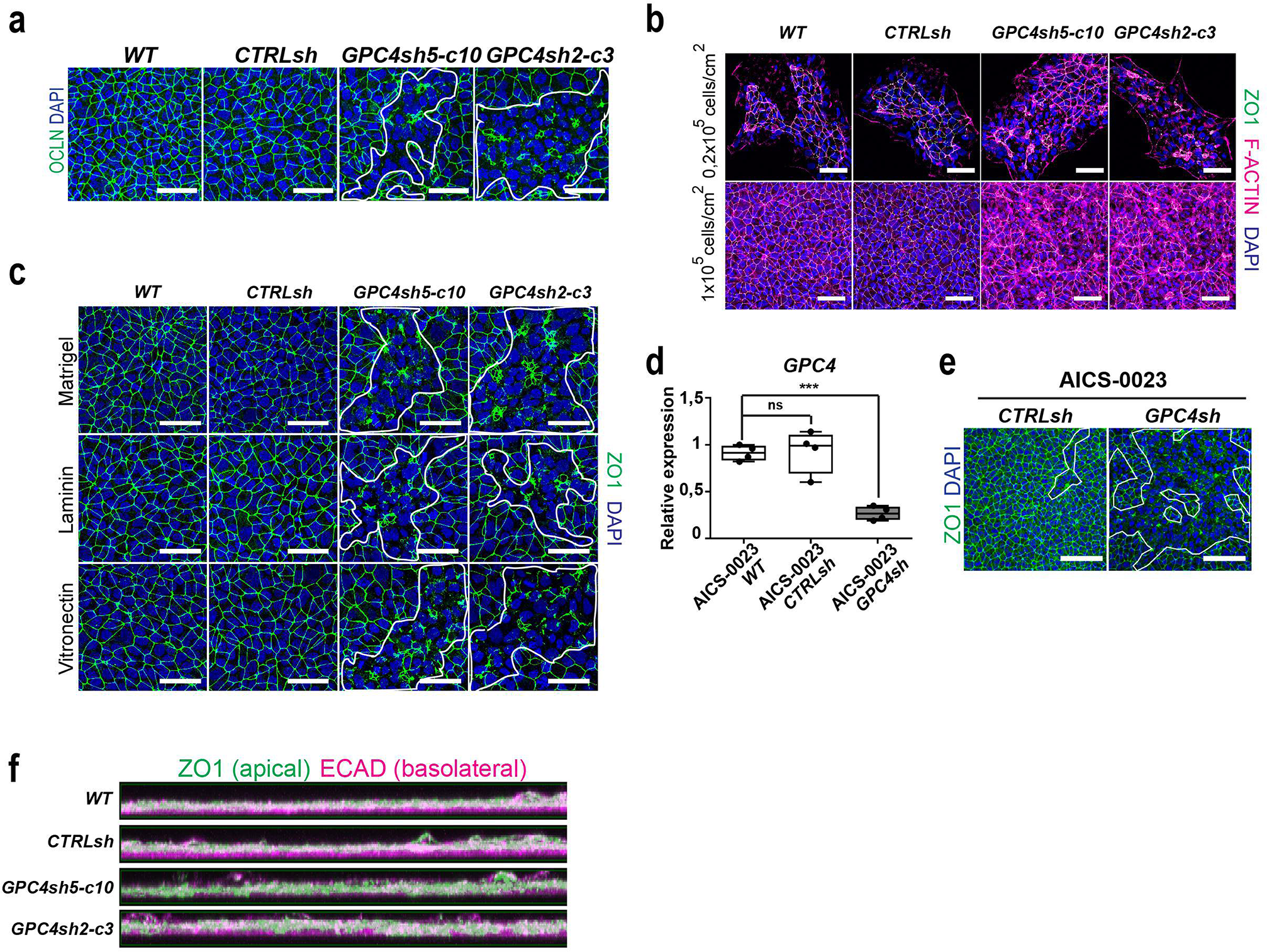
Down-regulation of GPC4 affects hiPSC epithelial integrity by disrupting TJs. **(a)** Immunofluorescence analysis of OCLN (green) in WT, CTRLsh, GPC4sh5-c10 and GPC4sh2-c3 029 hiPSCs showing that GPC4sh hiPSCs display extensive zones of disrupted TJs. Scale bar: 50 μm. **(b)** Immunofluorescence of F-ACTIN (magenta), ZO1 (green) and DAPI (blue) in WT, CTRLsh, GPC4sh5-c10 and GPC4sh2-c3 029 hiPSCs cultured at density of 0,2×10^5^ cells/cm^2^, and 1×10^5^ cells/cm^2^; showing that disruption of TJs observed in GPC4sh hiPSCs do not depend on cell density. Scale bar: 50 μm. **(c)** Immunofluorescence analysis of ZO1 (green) and DAPI (blue) in WT, CTRLsh, GPC4sh5-c10 and GPC4sh2-c3 029 hiPSCs cultured on Matrigel, Laminin and Vitronectin; showing that disruption of TJs observed in GPC4sh hiPSCs can be reproduced using different ECM. Scale bar: 50 μm. **(d)** RT-qPCR analyses of *GPC4* transcript levels in WT, CTRLsh and GPC4sh AICS-0023 hiPSC lines showing a down-regulation of *GPC4* expression in GPC4sh hiPSCs. Transcript levels were normalized to the mean of WT ΔCt, data are represented as median ± SD, n=4. **(e)** Immunofluorescence analysis of ZO1 (green) and DAPI (blue) in WT, CTRLsh and GPC4sh AICS-0023 hiPSC showing that GPC4sh hiPSCs display extensive zones of disrupted TJs. Scale bar: 50 μm. **(f)** Immunofluorescence analysis of ZO1 (green) and E-CAD (magenta) in WT, CTRLsh, GPC4sh5-c10 and GPC4sh2-c3 029 hiPSCs. Pictures are presented as lateral cell view. Statistical analysis for the overall figure: (b) one-way ANOVA followed by Dunnett’s multiple comparison test. P values: (***) < 0.001, (**) < 0.01, (*) < 0.05.

**Extended Data Fig. 2-1.**
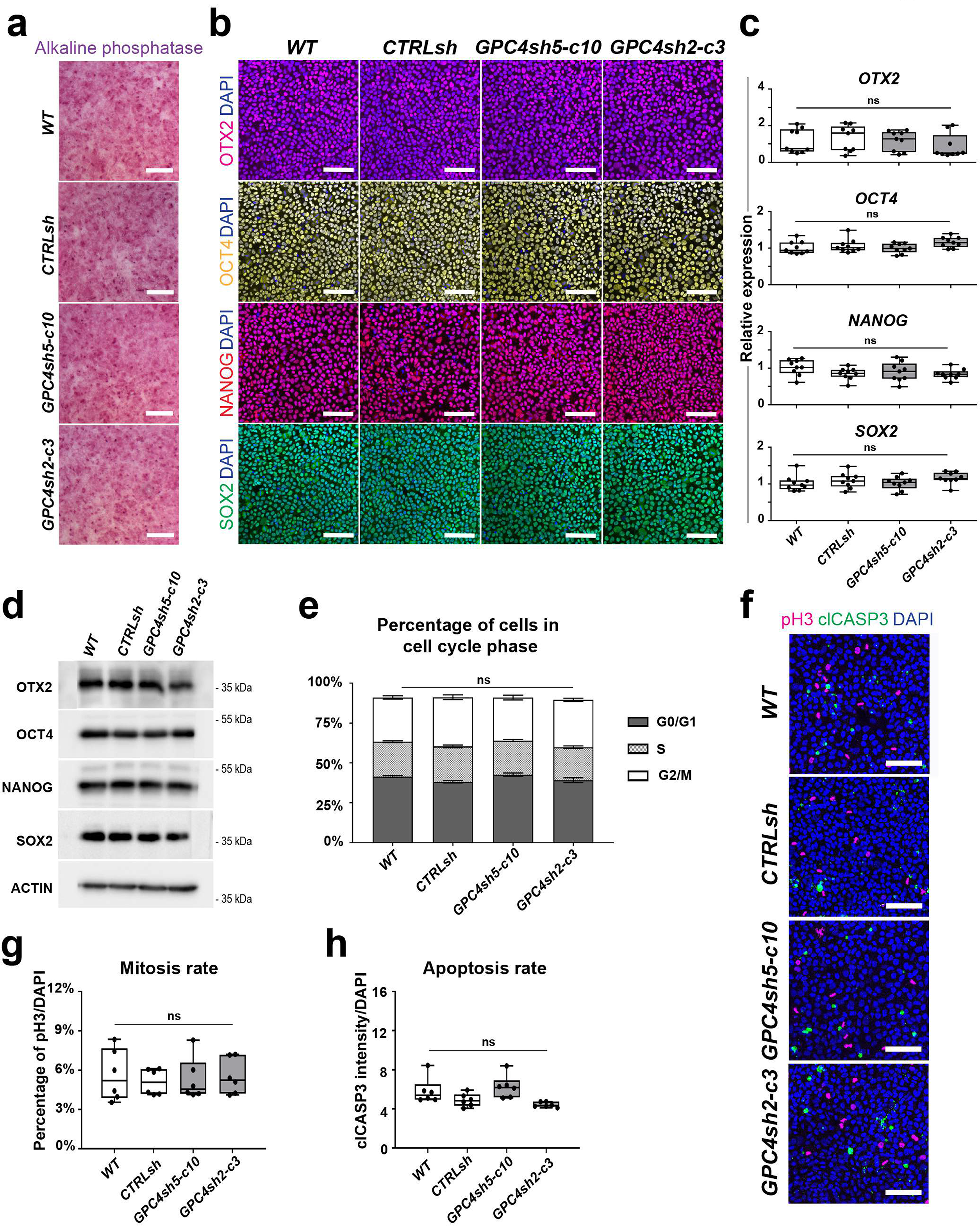
Disruption of epithelial integrity does not affect stemness properties of hiPSCs but enhances differentiation potential. **(a)** Alkaline phosphatase staining of WT, CTRLsh, GPC4sh5-c10 and GPC4sh2-c3 undifferentiated 029 hiPSCs. Scale bar: 100 μm. **(b)** Immunofluorescence analysis of the Epiblast marker OTX2 (magenta) and the pluripotency genes OCT4 (yellow), NANOG (red) and SOX2 (green) in WT, CTRLsh, GPC4sh5-c10 and GPC4sh2-c3 undifferentiated 029 hiPSCs. Scale bar: 100 μm. **(c)** Transcript levels of the Epiblast marker *OTX2* and the pluripotency genes *OCT4*, *NANOG* and *SOX2* were analyzed by RT-qPCR in WT, CTRLsh, GPC4sh5-c10 and GPC4sh2-c3 029 hiPSCs at the undifferentiated stage. Data were normalized to the mean of WT ΔCt. Results show no differences between WT, CTRLsh, GPC4sh5-c10 and GPC4sh2-c3 029 hiPSCs. Box plots represent the median with interquartile range, the errors bars indicate min and max values, n = 9. **(d)** Total protein extract of WT, CTRLsh, GPC4sh5-c10 and GPC4sh2-c3 029 hiPSCs at the undifferentiated stage were analyzed by western-blot with OTX2, OCT4, NANOG, and SOX2 antibodies. Note similar protein levels in all hiPSC lines. ACTIN protein levels were used as loading control. **(e)** Quantification of the percentage of cell in the phases G0/G1, S or G2/M of the cell cycle. Results show no differences between WT, CTRLsh, GPC4sh5-c10 and GPC4sh2-c3 029 hiPSCs. Box plots represent the median with interquartile range, the errors bars indicate min and max values, n=6. **(f)** Immunofluorescence analysis of pH3 (magenta) and cl-CASP3 (green) in undifferentiated WT, CTRLsh, GPC4sh5-c10 and GPC4sh2-c3 29 hiPSCs. Scale bar: 100 μm. **(g)** Percentages of pH3 positive cells were quantified from staining shown in (e). Data are represented as median ± SD. n = 3. **(h)** Global intensities of cl-CASP3 were quantified from staining shown in (e). Box plots represent the median with interquartile range, the errors bars indicate min and max values, n = 3. Statistical analysis for the overall figure: (c, e, g, h) one-way ANOVA followed by Dunnett’s multiple comparison test. P values: (***) < 0.001, (**) < 0.01, (*) < 0.05.

**Extended Data Fig. 2-2.**
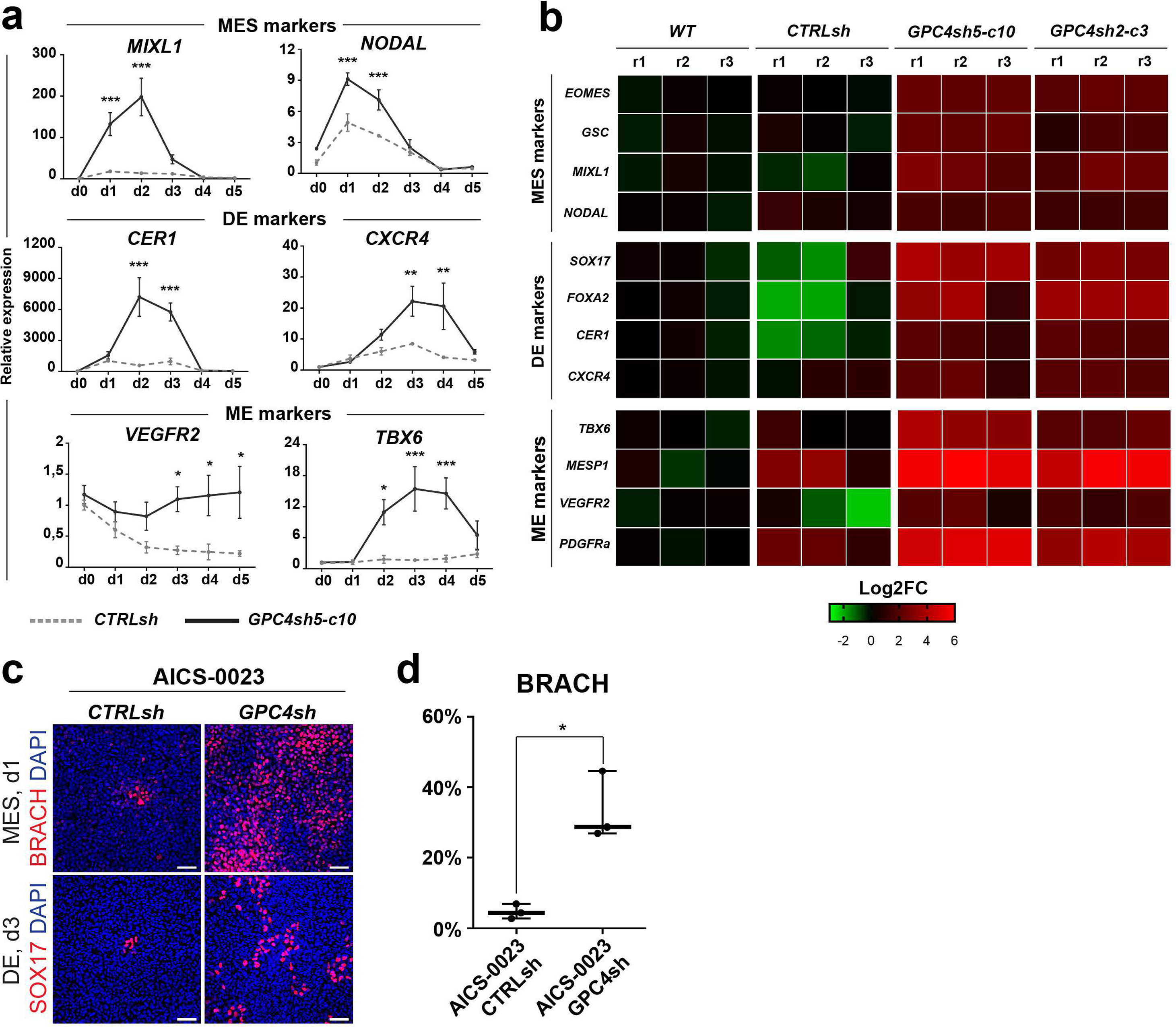
Disruption of epithelial integrity enhances differentiation potential. **(a)** CTRLsh and GPC4sh5-c10 029 hiPSCs were differentiated for 5 days into DE or ME and transcript levels of MES, DE or ME specific markers were subsequently analyzed by RT-qPCR. Top panel: MES markers (*MIXL1* and *NODAL*), middle panel: DE markers (*CER1* and *CXCR4*) and bottom panel: ME markers (*VEGFR2* and *TBX6*). Data were normalized to the d0 of CTRLsh and represented as mean ± SEM, n = 3. **(b)** WT, CTRLsh, GPC4sh5-c10 and GPC4sh2-c3 029 hiPSCs were differentiated for 1 day into MES, 3 days into DE or 3 days into ME, and transcript levels of lineage specific markers were analyzed by RT-qPCR. Top panel: MES markers (*EOMES*, *GSC*, *MIXL1* and *NODAL*), middle panel: DE markers (*SOX17*, *FOXA2*, *CER1* and *CXCR4*) and bottom panel: ME markers (*TBX6*, *MESP1*, *VEGFR2* and *PDGFRa*). Transcript levels were normalized to the mean of WT ΔCt, data are represented as heatmap of Log2FC, n = 3. **(c)** Immunofluorescence of the MES marker, BRACH, and the DE marker, SOX17, in CTRLsh and GPC4s AICS-0023 hiPSCs differentiated for 1 day into MES and 3 days into DE. Scale bar: 100 μm **(d)** Percentages of BRACH positive cells were quantified from staining shown in (c). Box plots represent the median with interquartile range, the errors bars indicate min and max values, n = 3. For the overall figure: one-way Anova statistical analysis: (***) < 0.001, (**) < 0.01, (*) < 0.05. Statistical analysis for the overall figure: (d) one-way ANOVA or (a) two-way ANOVA followed by Sidak’s multiple comparison test. P values: (***) < 0.001, (**) < 0.01, (*) < 0.05.

**Extended Data Fig. 3.**
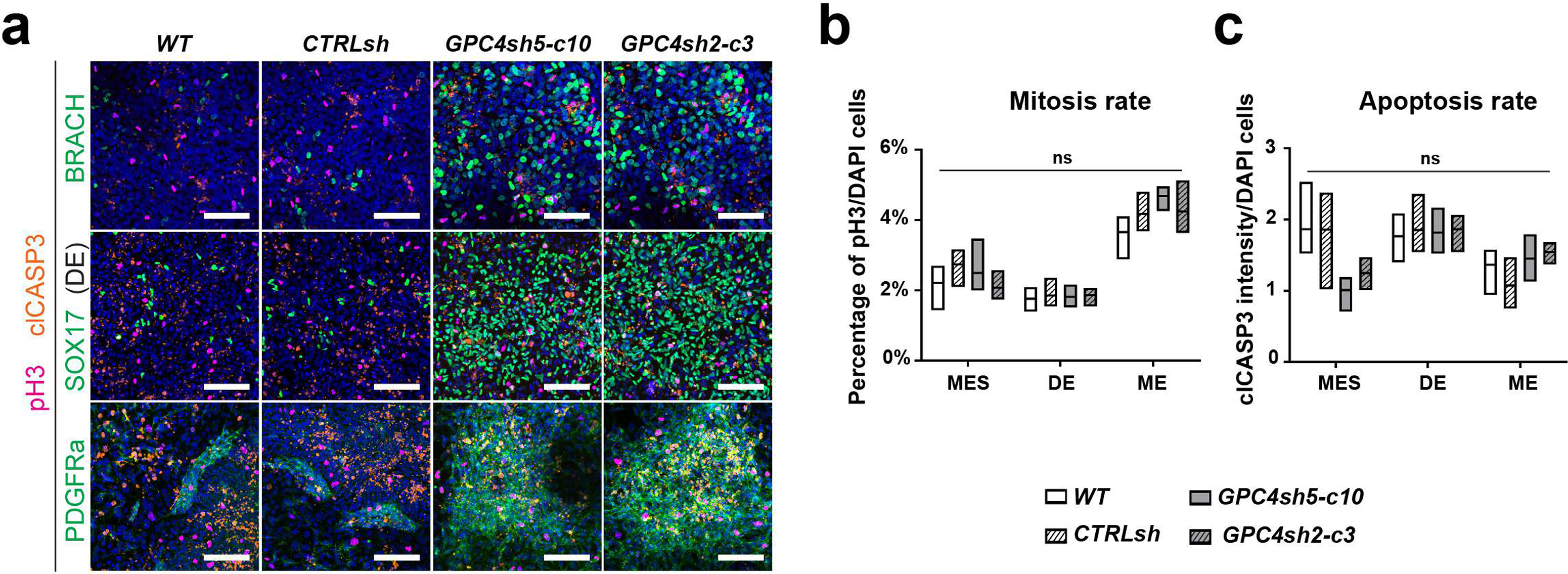
Analysis of cell differentiation properties in GPC4sh hiPSCs. **(a)** Immunofluorescence analysis of pH3 (magenta), cl-CASP3 (orange) with BRACH (green, PS), SOX17 (green, DE) or PDGFRa (green, ME) in WT, CTRLsh, GPC4sh5-c10 and GPC4sh2-c3 029 hiPSCs differentiated for 1 day into PS, 3 days into DE or 3 days into ME. Scale bar: 100 μm. **(b)** Percentages of pH3 positive cells were quantified from staining shown in (d). Box plots represent the median with interquartile range, the errors bars indicate min and max values, n = 3. **(c)** Global intensities of cl-CASP3 were quantified from staining shown in (d). Box plots represent the median with interquartile range, the errors bars indicate min and max values, n = 3. Statistical analysis for the overall figure: (a, e, f) two-way ANOVA followed by Sidak’s (a) or Tukey’s (e, f) multiple comparison tests. P values: (***) < 0.001, (**) < 0.01, (*) < 0.05.

**Extended Data Fig. 5.**
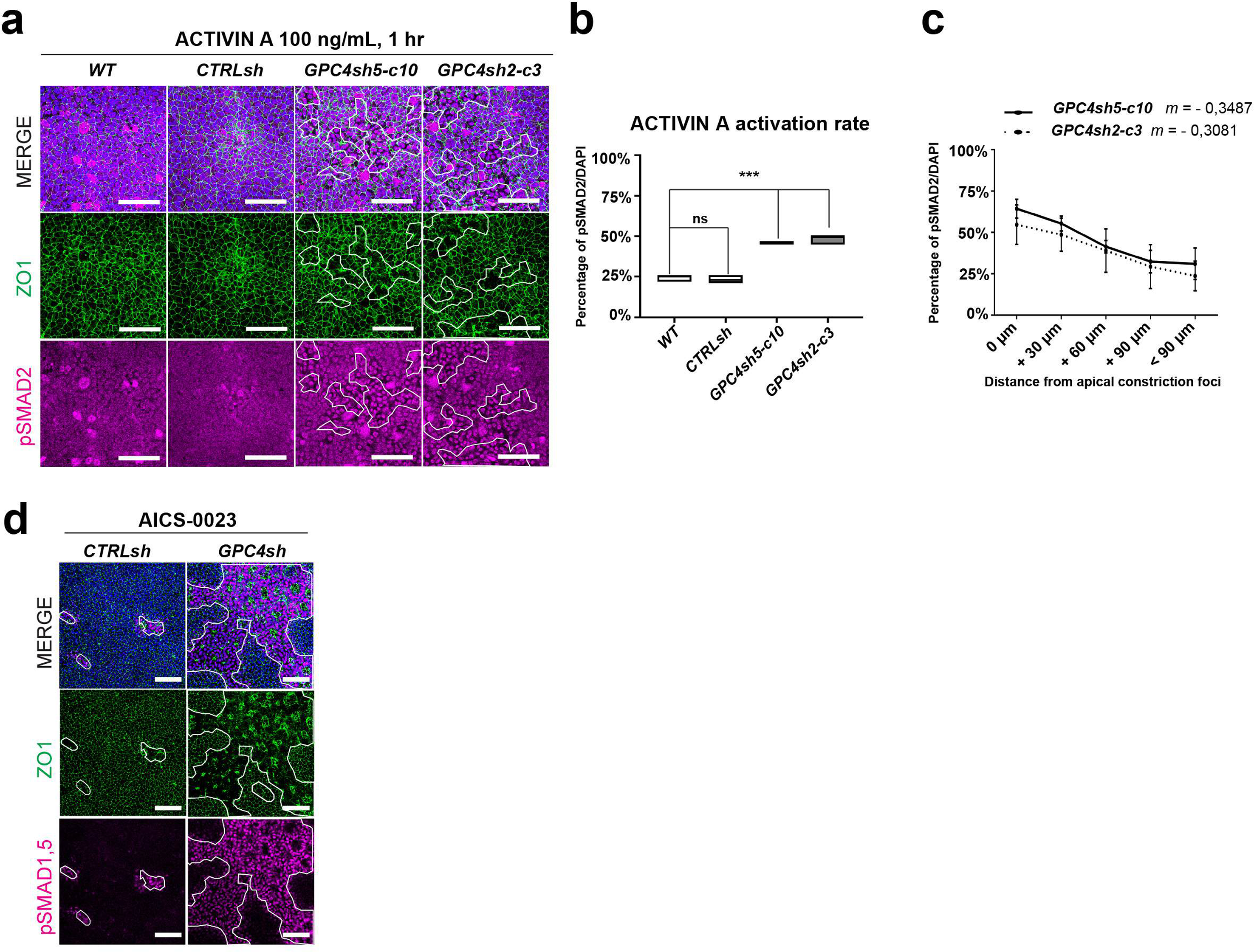
Epithelial integrity regulates TGF-β signaling efficiency and patterning. **(a)** Immunofluorescence analysis of pSMAD2 (magenta) and ZO1 (green) of WT, CTRLsh, GPC4sh5-c10, GPC4sh2-c3 029 hiPSCs stimulated for 1 hour with 100 ng/mL of ACTIVIN A. Dashed areas highlight the extent of zones of disrupted TJs, scale bar: 50 μm. **(b)** Percentages of pSMAD2 positive cells were quantified from staining shown in (a). Box plots represent the median with interquartile range, the errors bars indicate min and max values, n = 3. **(c)** Percentages of pSMAD2 ^S465/S467^ positive cells as a function of distance from the zone of disrupted TJs were quantified in GPC4sh5-c10 and GPC4sh2-c3 029 hiPSCs from staining shown in (A). 0 μm indicates cells located within the zones of disrupted TJs, + 30 μm, + 60 μm, + 90 μm and < + 90 μm indicates cells located at 0 to 30µm, to 60µm, 60 to 90µm or more than 90µm away from the zones of disrupted TJs, respectively. Data are represented as mean ± SD, n = 3. **(d)** Immunofluorescence analysis of pSMAD1,5 (magenta) and ZO1 (green) in cultures of CTRLsh and GPC4sh AICS-0023 hiPSCs stimulated for 1 hour with 50 μg/mL of BMP4. Dashed areas highlight the extent of zones of disrupted TJs. Scale bar: 100 μm.

**Extended Data Fig. 6.**
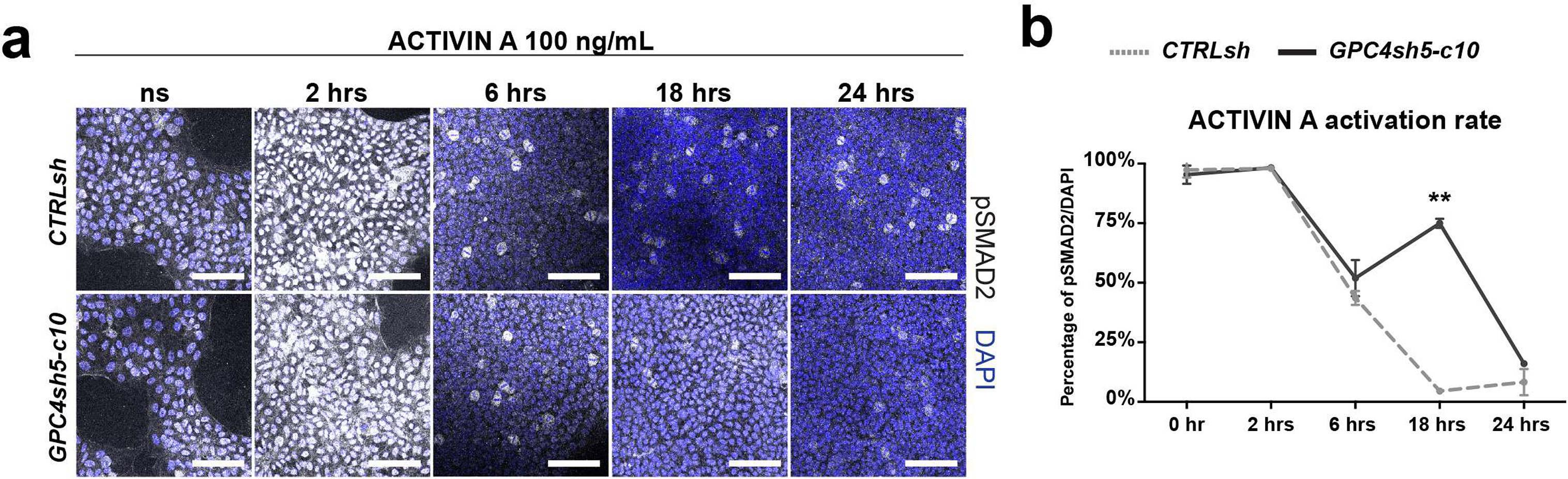
Disruption of epithelial integrity prolongs TGF-β signaling activation. **(a)** Immunofluorescence analysis of pSMAD2 (grey) in low densities cultures of CTRLsh and GPC4sh5-c10 029 hiPSCs stimulated for 2, 6, 18 and 24 hours with 100 ng/mL of ACTIVIN A. Scale bar: 100 μm. **(b)** Percentages of pSMAD2 expressing cells were quantified from staining shown in (f). Data are represented as mean ± SEM, n = 2. Statistical analysis for the overall figure: (c) linear regression analysis, (b) one-way ANOVA followed by Dunnett’s multiple comparison test, (g) two-way ANOVA followed by Sidak’s multiple comparison test. P values: (***) < 0.001, (**) < 0.01, (*) < 0.05.

